# Real-Time Selective Sequencing with RUBRIC: Read Until with Basecall and Reference-Informed Criteria

**DOI:** 10.1101/460014

**Authors:** Harrison S. Edwards, Raga Krishnakumar, Anupama Sinha, Sara W. Bird, Kamlesh D. Patel, Michael S. Bartsch

## Abstract

The Oxford MinION, the first commercial nanopore sequencer, is also the first to implement molecule-by-molecule real-time selective sequencing or “Read Until”. As DNA transits a MinION nanopore, real-time pore current data can be accessed and analyzed to provide active feedback to that pore. Fragments of interest are sequenced by default, while DNA deemed non-informative is rejected by reversing the pore bias to eject the strand, providing a novel means of background depletion and/or target enrichment. In contrast to the previously published pattern-matching Read Until approach, our RUBRIC method is the first example of real-time selective sequencing where on-line basecalling enables alignment against conventional nucleic acid references to provide the basis for sequence/reject decisions. We evaluate RUBRIC performance across a range of optimizable parameters, apply it to mixed human/bacteria and CRISPR/Cas9-cut samples, and present a generalized model for estimating real-time selection performance as a function of sample composition and computing configuration.

## Background

The Oxford Nanopore Technologies (ONT) MinION sequencer represents a significant paradigm shift in the reach, applicability, and capability of nucleic acid sequencing technology^1^. Combining a portable form factor, simple library prep, long-read capability (kb to Mb)^2^, direct RNA sequencing^3^, and real-time data output, the MinION has been variously applied to forensic genotyping^4^, bacterial typing^5^, plant biology^6^, food safety^7^, environmental metagenomics^8,9^, cancer research^10,11^, antibiotic resistance studies^12,13^, and de novo genome assembly^14–16^. The small operational and logistical footprint of the MinION, combined with its real-time capabilities^17^, make it uniquely suited to diagnostics and surveillance in clinical and field-forward settings, where the MinION has already been applied to assay Ebola^18,19^, Zika^20^, tuberculosis^21^, and other pathogens^22–25^.

Despite these successes, nanopore sequencing-based diagnostics still face the “needle in a haystack” problem of obtaining sufficient coverage of low-abundance target from a high-abundance background (e.g., pathogen/host, cancer/nontumor) sample^26^. While bacterial culture provides enriched quantities of genetic material in some applications^27^, culture-independent molecular biology-based target enrichment and background depletion methods^28^ including amplification^29^ and hybridization capture approaches^30^ are increasingly being adapted for use in library preparation to yield “targeted” or “selective” sequencing^31,32^. Nearly all such methods require *a priori* knowledge to guide the design of the target-sequence-specific primers, baits, or probes required for selection.

Unique to the Oxford MinION, *real-time selective sequencing* was first introduced by Loose and colleagues in 2016^33^, offering a promising alternative to these molecular biology-based enrichment approaches. Dubbed “Read Until”, the method capitalizes on the real-time data output and discretely addressable nanopore architecture of the MinION to enable selection of individual DNA molecules. Read Until makes it possible to preview the real-time data associated with DNA traversing a given nanopore, and if it fails to meet some user-defined selection criteria, reject that read by reversing the pore bias and physically ejecting the DNA (i.e., “unblocking” the pore).

DNA meeting the criteria sequences to completion as usual, with selection producing a net enrichment of target versus non-target reads in the final sequence pool. Read Until sequence-based selection has no clear precedent in the literature, the closest analogs being size-based^34^ and methylation-based^35^ DNA sorting in nanochannels, while most “single-molecule sorting” methods principally consist of surface immobilization coupled with molecular-resolution fluorescence imaging^36^.

In the original Read Until implementation, Loose applied a dynamic time warping (DTW) algorithm to pattern-match the live current trace “squiggle” output by the MinKNOW sequencing software against a reference squiggle synthesized from the (ACGT) target sequence of interest^33^. The method was successfully executed at a time when the MinION sequencing rate was 70 bases/s (it is now 450 bases/s) using a 22-core server to select for 5 kb portions of lambda DNA and to normalize coverage among 2 kb amplicons. Subsequent work developed a statistical model for optimizing DTW selection^37^. Here we introduce a new implementation of real-time selective sequencing based on Loose’s original framework: Read-Until with Basecall and Reference-Informed Criteria (RUBRIC). Rather than pattern-matching event traces, RUBRIC relies on real-time basecalling and alignment to conventional ACGT-type reference sequences, providing significant benefits to speed, scalability, and operational flexibility. Moreover, RUBRIC is specifically designed to function with the more modest computing resources typical of portable or point-of-need MinION-based activities rather than high-end multiprocessor workstations or cluster computing platforms. In addition to characterizing the operation of the RUBRIC architecture for a series of proof-of-concept experiments, we also propose a predictive model evaluating the likely limits of real-time selection performance generally across a range of potential sample types and use cases.

## Methods

### RUBRIC Implementation & Operation

Figure 1 shows the RUBRIC real-time selection architecture, implemented with off-the-shelf, ethernet-linked laptop and desktop PCs, while Table 1 summarizes all RUBRIC experiments discussed here. Built upon the original Read Until sample code provided by Loose^33^, RUBRIC integrates ONT’s Nanonet basecaller (v2.0.0, included with the RUBRIC code as noted below) and replaces DTW-based target pattern-matching with sequence-based alignment using LAST (rev 759)^38^. For each sequencing experiment, initial MinKNOW calibration and multiplex scans were performed, MinKNOW sequencing was initiated, and RUBRIC scripts were then started on the desktop PC. Depicted in Figure 1, the general RUBRIC control flow consisted of receiving batches of read events from the Read Until Event Sampler, formatting those events for basecalling by Nanonet, aligning the results against a desired target reference sequence with LAST, and parsing its output to make skip/sequence determinations which were then communicated to MinKNOW via the Read Until API. LAST arguments used in the RUBRIC selection process are shown in Table 1. For all experiments, the Event Sampler was set to ignore the first 100 (typically lower fidelity^39^) events of each processed read and then transmit an “evaluation window” comprising the next 300 events (600 for run G, see Table 1) as the input to the RUBRIC selection process. During all experiments, the RUBRIC scripts logged relevant Event Sampler read information for method improvement and downstream reconciliation with offline Albacore basecall and BWA alignment results.

**Figure 1:**
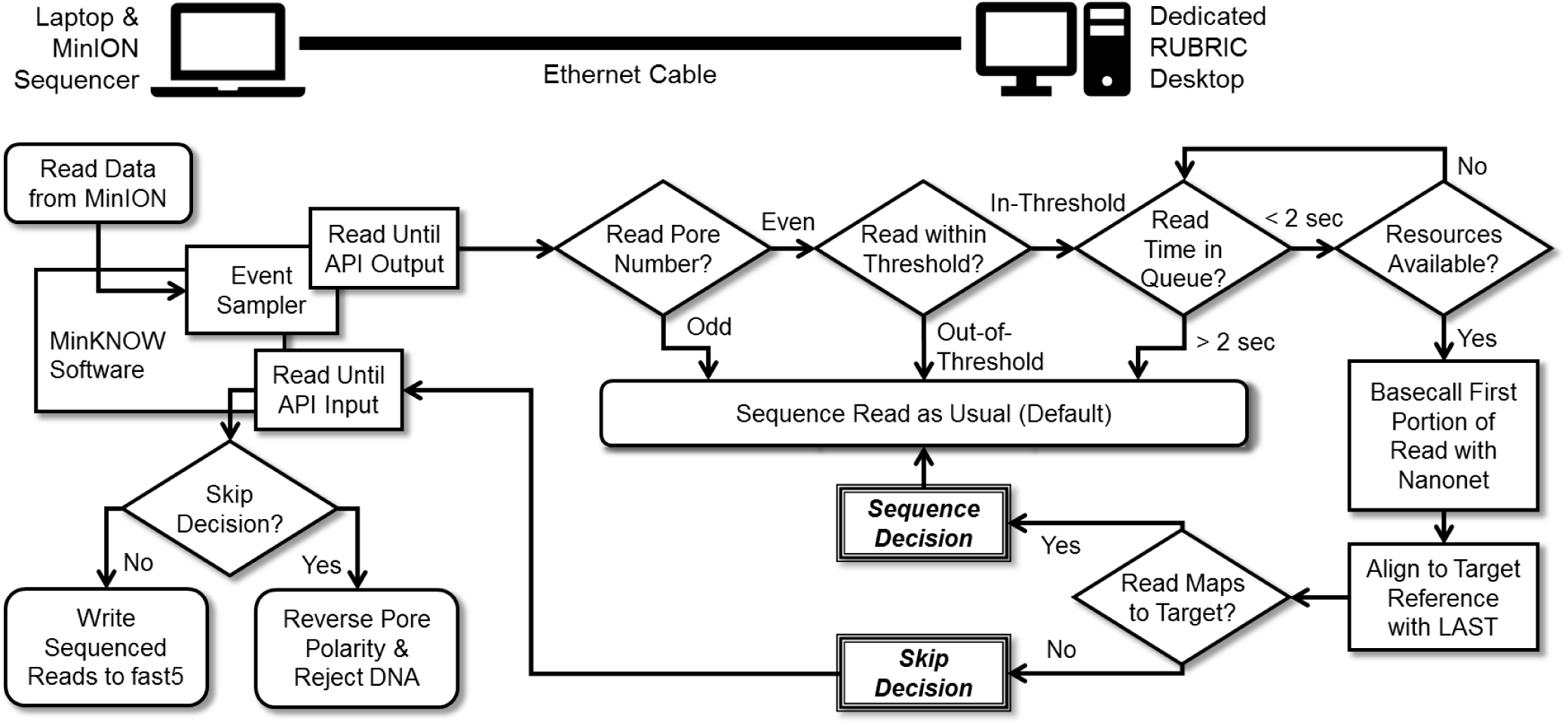
Schematic of the RUBRIC workflow illustrating the division of computational effort between two garden-variety PCs: a laptop that runs the MinION sequencer and its MinKNOW software interfaced through the Read Until API (via ethernet) to a desktop system that performs the key RUBRIC operations of pre-screening reads for admission to the decision process, basecalling and aligning reads to nucleic acid target reference(s) in real-time, and communicating any resulting skip/reject decisions back to MinKNOW.

**Table 1:**
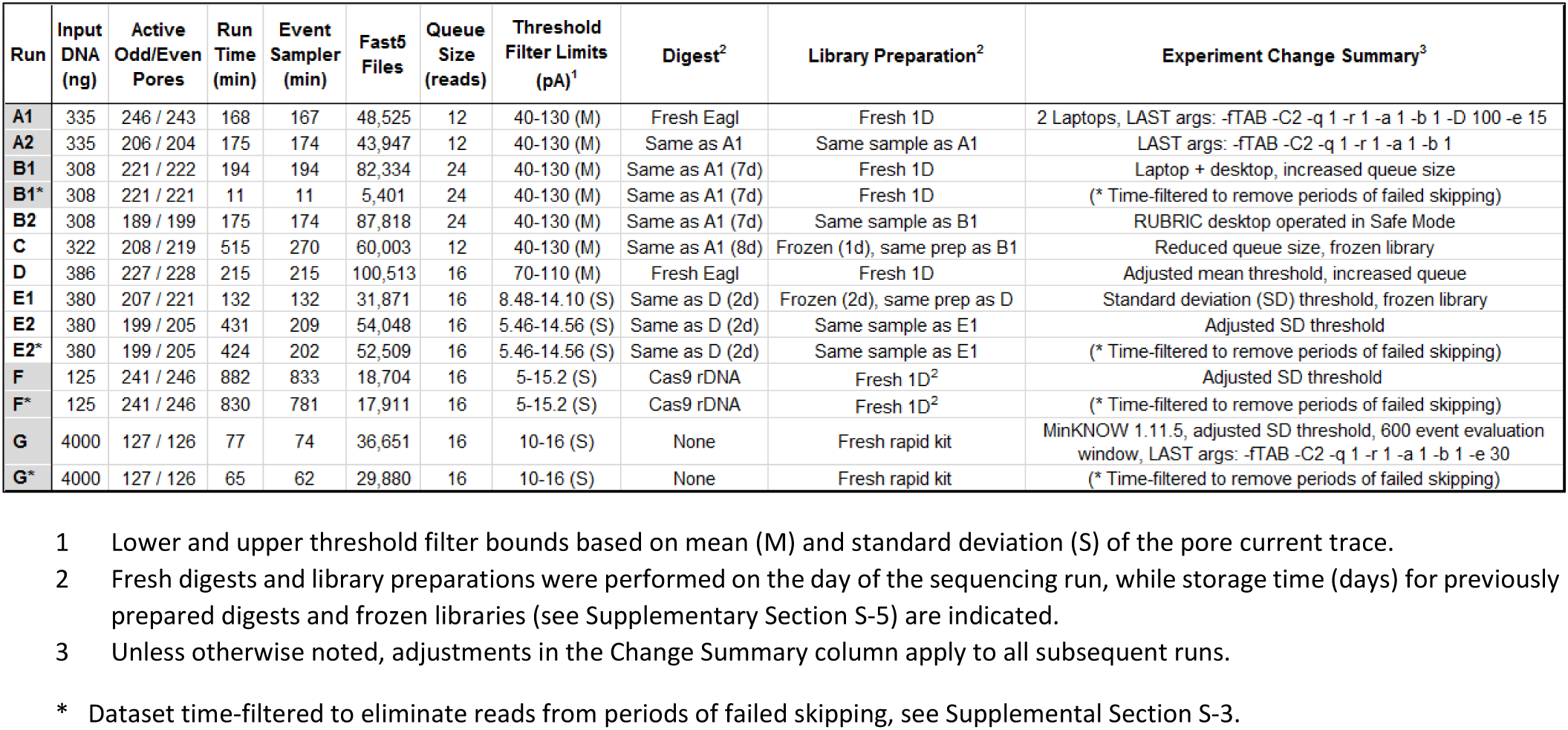
Summary of RUBRIC experiments and parametric variations for preliminary lambda DNA experiments A1-B1, mainline EagI-digested Lambda DNA experiments B2-E2, and example use case experiments F and G in which Cas9-cut rDNA was selected from E. coli gDNA and E. coli gDNA was selected from human gDNA, respectively.

Despite processing only a short initial portion of each read (∼150 bases from 300 events), successfully implementing RUBRIC with garden-variety PCs necessitated careful conservation of limited computing resources. In addition to running RUBRIC on a dedicated desktop machine, Figure 1 illustrates the additional steps that were taken to control the volume and optimize the relevance of reads admitted to the RUBRIC decision process. First, in all experiments detailed here, RUBRIC selection was applied only to even-numbered pores, while odd pores were allowed to sequence normally, providing an internal control. Second, a threshold filter was implemented by quickly computing the mean or standard deviation (Supplementary Section S-2) of pore current for the evaluation window, and on that basis, excluding from selection reads that were empirically determined to be unlikely to yield mappable fast5 sequence files. Lastly, a queue was implemented to: 1) constrain the number of event traces passed to RUBRIC at a given time to avoid overwhelming available computing resources and 2) screen reads that spent too long in the queue from entering the decision process. Queue size varied between 12 and 24 reads (Table 1), but in all experiments, reads spending more than 2 seconds in the queue were deemed too old for a timely decision to be rendered, and therefore bypassed selection. As Figure 1 indicates, during the RUBRIC development and characterization process, the default for any reads not admitted to the selection process (i.e., odd, out-of-threshold, timeout, and otherwise “undecided” reads) and for reads receiving an affirmative “sequence” decision was to sequence as usual. Only reads receiving a “skip” decision resulting in ejection by pore polarity reversal (unblocking) were not sequenced by default.

### Software & Computing Architecture

After a preliminary experimental iteration using two laptop PCs (Table 1, runs A1-A2), the final and preferred RUBRIC sequencing setup (Figure 1) consisted of an off-the-shelf HP Elitebook 820 G3 laptop with 4 cores (Intel^®^ Core™ i7-6500U CPU @ 2.5 GHz, 16 GB RAM, Samsung MZNLN512HCJH-000H1 477GB SCSI SSD) connected by USB to a MinION Mk1B sequencer and by 2-foot Cat-5e Ethernet cable to a Dell Optiplex 9020 desktop with 8 cores (Intel^®^ Core™ i7-4790 CPU @ 3.6 GHz, 16 GB RAM, Samsung 850 2TB SCSI SSD). Oxford MinKNOW version 1.6.11 sequencing software was run on the laptop for all experiments other than run G (v1.11.5), while the desktop system provided the additional computing power needed to implement RUBRIC real-time basecalling, alignment, and selection functions concurrently with sequencing. No other computing resources were used within the RUBRIC control loop. RUBRIC software communicated with MinKNOW’s Event Sampler via the Read Until API (v1) to acquire event data and provide rejection instructions in real time. Both computers operated in Windows 10, and the desktop was placed into Safe Mode during runs to prevent CPU usage by background processes and services. After sequencing, all data were basecalled offline using Albacore v1.2.6 (v2.2.4 for run G) and post-run alignment was performed using BWA v0.7.12-r1039 (with ‘mem -x pacbio’ arguments) on Sandia’s Biota computing cluster. While BWA was used for offline alignment and classification of output MinION reads, LAST was selected for use inside the RUBRIC control loop due to its speed and the comparative ease of integrating it into the real-time workflow. Downstream data analysis and visualization were performed using custom Python scripts (pandas, numpy, matplotlib, seaborn), custom R scripts, and Microsoft Excel.

### Sample Preparation & Experimental Variations

#### Lambda DNA Experiments

To provide a test case for RUBRIC selection, lambda-phage DNA (cat # N3011S, New England Biolabs (NEB), Ipswich, MA) was digested using the EagI enzyme (NEB, cat # R3505S) to produce three large DNA fragments of roughly similar size (20 kb, 17 kb, and 12 kb). Digestion was performed per NEB protocol in a 50 µL reaction, and the product was purified using phenol:chlororform. The 17 kb fragment was chosen as the target for RUBRIC selection, while reads not matching its sequence were skipped. For all lambda DNA experiments (A1-E2 in Table 1), digested samples were prepared using ONT’s 1D ligation kit (SQK-LSK108) and loaded into SpotON flow cells (FLO-MIN107, used for all experiments in this article) using methods described in the kit’s accompanying protocol. DNA concentrations were measured using a Qubit Fluorimeter (Thermo Fisher, Waltham, MA).

Table 1 summarizes the progression of experimental parameter variations through sequential RUBRIC experiments, with letters differentiating experiments performed on different days and numbers indicating successive RUBRIC runs with the same loaded sample (but different RUBRIC settings) on a given day. Datasets indicated with an asterisk (*) have been time-filtered as explained in Supplementary Section S-3 to eliminate data from periods during which skip decisions failed to properly reject DNA. Experiments A1, A2, and B1 are included primarily for comparison, reflecting the earliest parametric iterations and system configurations, and are therefore not representative of typical RUBRIC performance. Accordingly, aggregate results distinguish between “mainline” results associated with the preferred RUBRIC system configuration (N=5, runs B2-E2), and the set of all lambda experiments (N=8, A1-E2). Non-lambda DNA runs F and G, described below, are preliminary proof-of-concept examples applying RUBRIC in use cases potentially relevant to pathogen diagnostics.

To summarize the variations tested for lambda DNA, runs A1 and A2, performed using two equivalent, Ethernet-coupled laptops, tested the effect of changing the settings of the LAST aligner used in the RUBRIC control loop. Experiment B1 used the same settings but implemented RUBRIC on ethernet-linked laptop and desktop machines, while B2 revealed the benefit of operating the RUBRIC-running desktop in Safe Mode. Experiment C used a previously prepared frozen library and reduced the queue size from 24 to 12. Experiment D increased the queue to 16 and adjusted the mean current-based threshold with a fresh digest and library prep. Experiment E1 implemented a standard deviation-based threshold for a frozen library, and experiment E2 further adjusted that threshold.

#### E. coli Ribosomal DNA Experiment

While long-fragment lambda DNA proof of concept experiments facilitated early RUBRIC optimization and troubleshooting efforts, we also performed preliminary experiments to assess the potential of RUBRIC selection in more realistic applications, specifically with an eye toward bacterial pathogen diagnostics. In experiment F, inspired by conventional bacterial ribotyping, guide RNAs were designed to target the 5’ end of the 16S and the 3’ end of the 23S ribosomal DNA (rDNA) loci of *E. coli* (Accession number: NC_000913) to excise the ∼5kb 16S-23S region of the rDNA locus. Single-molecule guide RNA (sgRNA) templates were generated by polymerase chain reaction (PCR) (16S primer 5’-M-TGGCTCAGATTGAACGCTGG-N-3’ and 23S primer 5’-M-CGCCCAAGAGTTCATATCGA-N-3’, where M=5’-GGATCCTAATACGACTCACTATAG-3’ and N=5’-GTTTTAGAGCTAGAA-3’) to yield a single chimeric template containing the crRNA, tracrRNA, and a T7 promoter sequence as described by Anders^40^. sgRNAs were transcribed *in vitro* using the TranscriptAid T7 High Yield Transcription Kit (Thermo Fisher, cat # K0441) according to manufacturer’s protocol. Guide RNAs were purified using MEGAclear Transcription Clean-Up Kit (Thermo Fisher/Ambion, cat # AM1908) according to manufacturer’s protocol and diluted to 300nM.

For the CRISPR/Cas9 digest, a 90 µL reaction was prepared by mixing 9 µL of 10X Cas9 Nuclease Reaction Buffer (NEB), 30 nM gRNA1 (targeting 16S region), 30 nM gRNA2 (targeting 23S region) and 30 nM SpyCas9 Nuclease (NEB, cat#M0386S). After a 15 min incubation to form the ribonucleoprotein complex, 10 µg of bacterial genomic DNA was added and the reaction incubated at 37 °C for 4 hours. 1 µL of proteinase K (Thermo Fisher, AM2548) was added and the reaction incubated at 65 °C for 15 minutes. DNA was purified using Agencourt AMPure XP beads (cat #A63881, Beckman-Coulter, Brea, CA) according to manufacturer’s protocol. Library preparation was performed per ONT protocol using the 1D^2^ ligation kit (SQK-LSK308), and RUBRIC targets were set to select for the 16S-23S rDNA sequences (NCBI).

#### Mixed Human/E. coli Experiment

The second example use case, experiment G, sought to select for 1% *E. coli* genomic DNA against a background of 99% human DNA (HeLa, NEB, cat# N4006S) in a sample mixed prior to library preparation. *Escherichia coli* K12 MG1655 (ATCC, Manassas, VA) culture was grown overnight in LB media at 37 °C with shaking at 250 rpm. 1 mL aliquots were spun down to make the bacterial pellet, and cells were lysed using Qiagen lysis buffer (Qiagen, Redwood City, CA) with added Proteinase K and RNase A (Thermo Fisher). The lysate mixture was incubated for 15-30 min at 50 °C. Pure genomic DNA was extracted using the phenol:chloroform extraction method. Briefly, one volume of phenol:chloroform:isoamyl alcohol (25:24:1) (Sigma-Aldrich, St. Louis, MO) was added to the lysate mixture and the samples were centrifuged at room temperature for 10 minutes at 16,000 × g. The upper aqueous phase was transferred to a fresh tube and the DNA was precipitated by the addition of 0.1 volumes 3 M sodium acetate (pH 5.0) and 2.5 volumes of 100% ethanol. The samples were stored at −20°C overnight to precipitate the DNA. The DNA was pelleted at 4 °C for 15-30 minutes at 16,000 × g and the DNA pellets were washed twice with 500 µL of 70% ethanol. The DNA pellets were dried at room temperature for 5-10 minutes and resuspended in nuclease free water, and library preparation was accomplished using a RAD004 rapid kit per ONT protocol. During RUBRIC operation, reads were LAST-aligned in real-time against the entire 4.6 Mb *E. coli* K12 genome (NCBI) as the selection target. As noted in Table 1, for experiment G the evaluation window was increased from 300 to 600 events to enable greater discrimination between bacterial and human sequence, and LAST stringency was reduced to capture as many rare target reads as possible.

## Results

### Data Flow Analysis & Lambda DNA Results

Figure 2 illustrates the detailed data flow analysis approach used to evaluate even pore RUBRIC selective sequencing performance in comparison to the internal control provided by non-selecting odd channels for representative lambda DNA experiment B2. Equivalent Sankey diagrams for all other experiments (and filtered datasets) are provided in Supplementary Figure S9 with results summarized in Supplementary Figure S1. Table 2 compares performance metrics for the runs.

**Figure 2:**
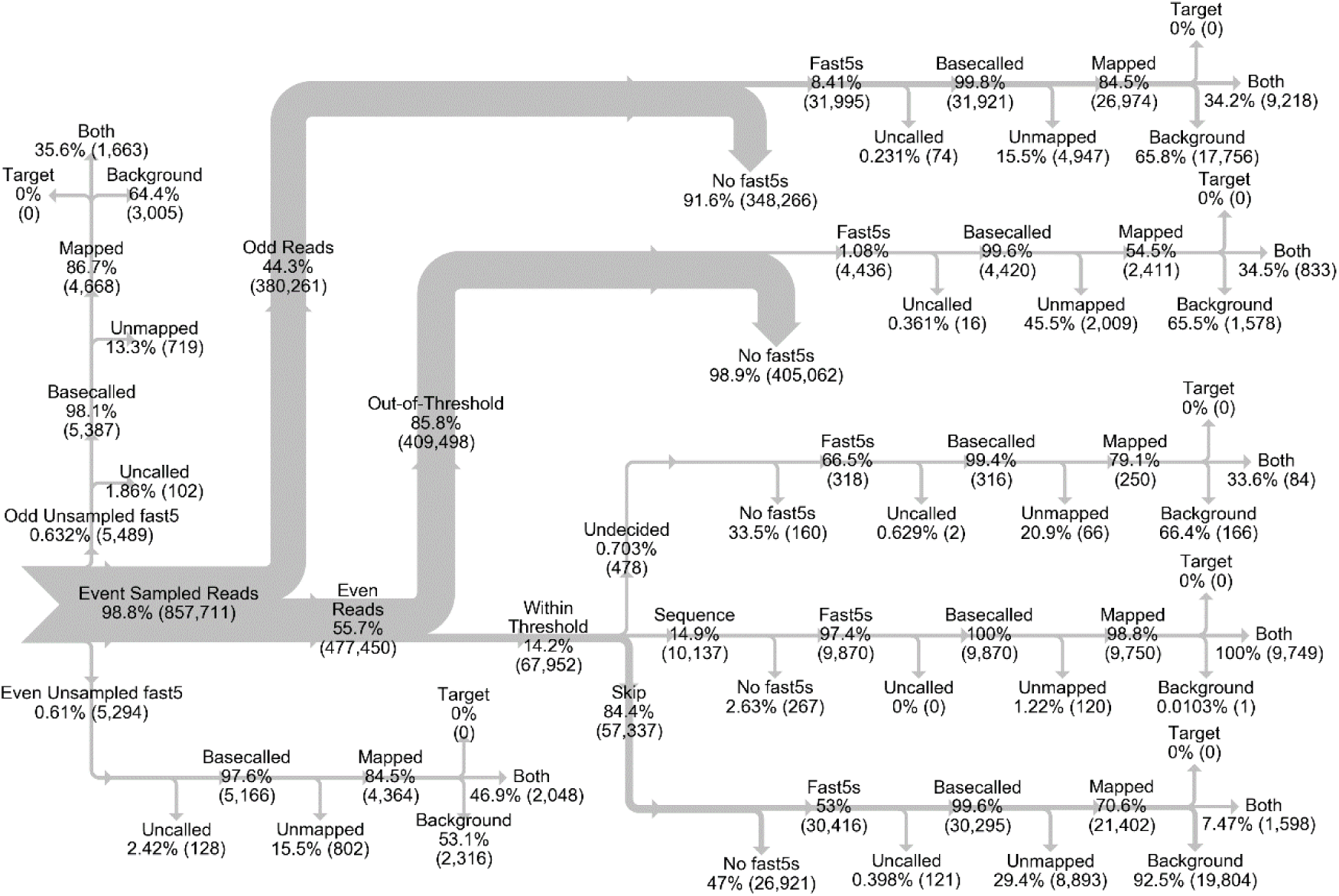
Sankey chart depicting read and fast5 sequence file data flow analysis for Experiment B2. Because the target lambda DNA fragment was a subset of the overall lambda (background) sequence, no reads mapped exclusively to the target, and therefore all correctly mapped target reads appear in the “both” category at the 3-pronged terminal ends of each chart branch. Undecided read counts shown here include both reads that timed-out of the decision process (> 2 seconds in the queue) and those that did not otherwise receive a decision.

**Table 2:**
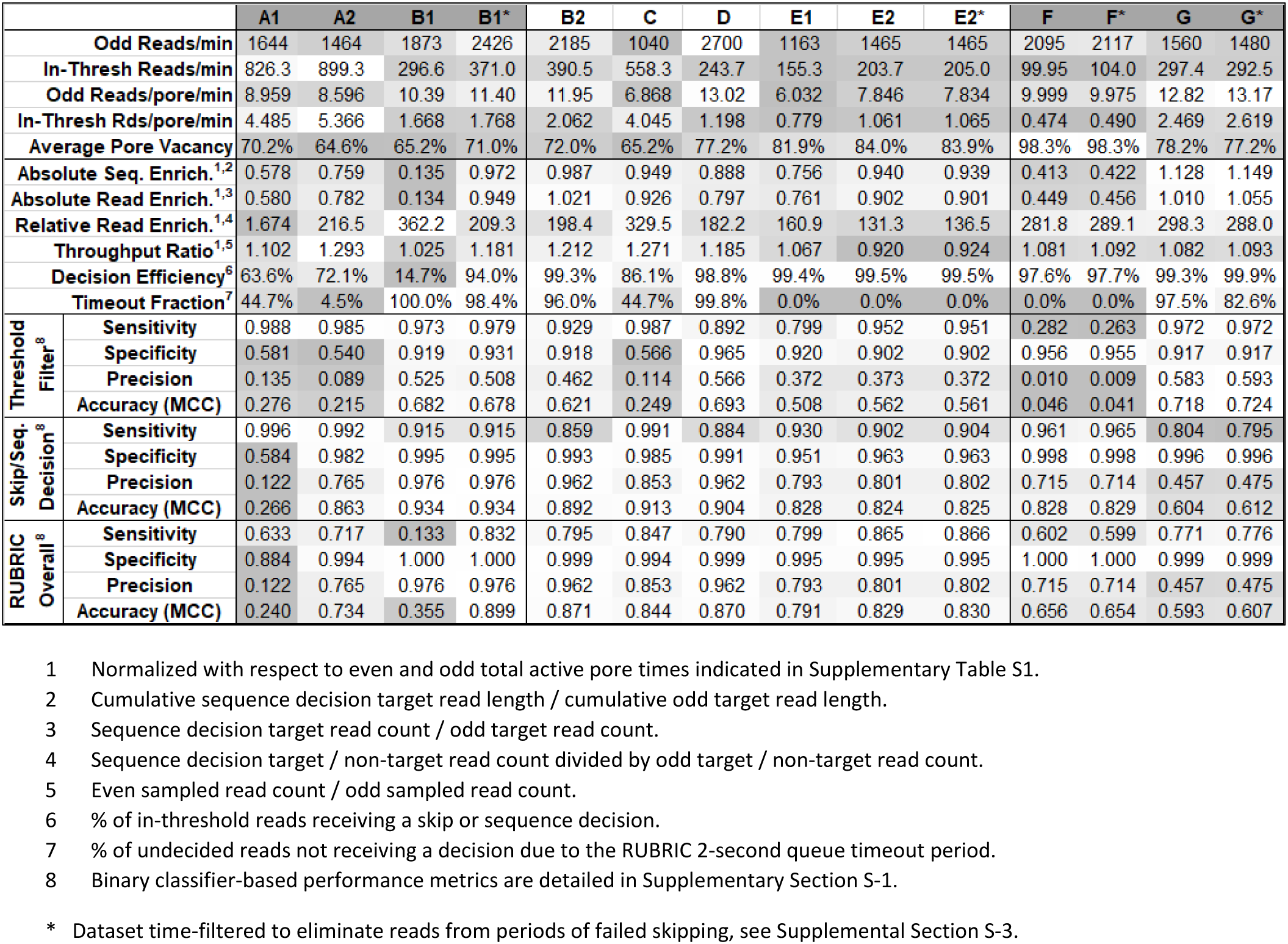
Performance metrics for RUBRIC selective sequencing experiments with shading indicating low (dark) and high (light) values within each row.

|  |  | A1 | A2 | B1 | B1* | B2 | C | D | E1 | E2 | E2* | F | F* | G | G* |
| --- | --- | --- | --- | --- | --- | --- | --- | --- | --- | --- | --- | --- | --- | --- | --- |
| Odd Reads/min |  | 1644 | 1464 | 1873 | 2426 | 2185 | 1040 | 2700 | 1163 | 1465 | 1465 | 2095 | 2117 | 1560 | 1480 |
| In-Thresh Reads/min |  | 826.3 | 899.3 | 296.6 | 371.0 | 390.5 | 558.3 | 243.7 | 155.3 | 203.7 | 205.0 | 99.95 | 104.0 | 297.4 | 292.5 |
| Odd Reads/pore/min |  | 8.959 | 8.596 | 10.39 | 11.40 | 11.95 | 6.868 | 13.02 | 6.032 | 7.846 | 7.834 | 9.999 | 9.975 | 12.82 | 13.17 |
| In-Thresh Rds/pore/min |  | 4.485 | 5.366 | 1.668 | 1.768 | 2.062 | 4.045 | 1.198 | 0.779 | 1.061 | 1.065 | 0.474 | 0.490 | 2.469 | 2.619 |
| Average Pore Vacancy |  | 70.2% | 64.6% | 65.2% | 71.0% | 72.0% | 65.2% | 77.2% | 81.9% | 84.0% | 83.9% | 98.3% | 98.3% | 78.2% | 77.2% |
| Absolute Seq. Enrich.<sup>1,2</sup> |  | 0.578 | 0.759 | 0.135 | 0.972 | 0.987 | 0.949 | 0.888 | 0.756 | 0.940 | 0.939 | 0.413 | 0.422 | 1.128 | 1.149 |
| Absolute Read Enrich.<sup>1,3</sup> |  | 0.580 | 0.782 | 0.134 | 0.949 | 1.021 | 0.926 | 0.797 | 0.761 | 0.902 | 0.901 | 0.449 | 0.456 | 1.010 | 1.055 |
| Relative Read Enrich.<sup>1,4</sup> |  | 1.674 | 216.5 | 362.2 | 209.3 | 198.4 | 329.5 | 182.2 | 160.9 | 131.3 | 136.5 | 281.8 | 289.1 | 298.3 | 288.0 |
| Throughput Ratio<sup>1,5</sup> |  | 1.102 | 1.293 | 1.025 | 1.181 | 1.212 | 1.271 | 1.185 | 1.067 | 0.920 | 0.924 | 1.081 | 1.092 | 1.082 | 1.093 |
| Decision Efficiency<sup>6</sup> |  | 63.6% | 72.1% | 14.7% | 94.0% | 99.3% | 86.1% | 98.8% | 99.4% | 99.5% | 99.5% | 97.6% | 97.7% | 99.3% | 99.9% |
| Timeout Fraction<sup>7</sup> |  | 44.7% | 4.5% | 100.0% | 98.4% | 96.0% | 44.7% | 99.8% | 0.0% | 0.0% | 0.0% | 0.0% | 0.0% | 97.5% | 82.6% |
| Threshold Filter<sup>8</sup> | Sensitivity | 0.988 | 0.985 | 0.973 | 0.979 | 0.929 | 0.987 | 0.892 | 0.799 | 0.952 | 0.951 | 0.282 | 0.263 | 0.972 | 0.972 |
|  | Specificity | 0.581 | 0.540 | 0.919 | 0.931 | 0.918 | 0.566 | 0.965 | 0.920 | 0.902 | 0.902 | 0.956 | 0.955 | 0.917 | 0.917 |
|  | Precision | 0.135 | 0.089 | 0.525 | 0.508 | 0.462 | 0.114 | 0.566 | 0.372 | 0.373 | 0.372 | 0.010 | 0.009 | 0.583 | 0.593 |
|  | Accuracy (MCC) | 0.276 | 0.215 | 0.682 | 0.678 | 0.621 | 0.249 | 0.693 | 0.508 | 0.562 | 0.561 | 0.046 | 0.041 | 0.718 | 0.724 |
| Skip/Seq. Decision<sup>8</sup> | Sensitivity | 0.996 | 0.992 | 0.915 | 0.915 | 0.859 | 0.991 | 0.884 | 0.930 | 0.902 | 0.904 | 0.961 | 0.965 | 0.804 | 0.795 |
|  | Specificity | 0.584 | 0.982 | 0.995 | 0.995 | 0.993 | 0.985 | 0.991 | 0.951 | 0.963 | 0.963 | 0.998 | 0.998 | 0.996 | 0.996 |
|  | Precision | 0.122 | 0.765 | 0.976 | 0.976 | 0.962 | 0.853 | 0.962 | 0.793 | 0.801 | 0.802 | 0.715 | 0.714 | 0.457 | 0.475 |
|  | Accuracy (MCC) | 0.266 | 0.863 | 0.934 | 0.934 | 0.892 | 0.913 | 0.904 | 0.828 | 0.824 | 0.825 | 0.828 | 0.829 | 0.604 | 0.612 |
| RUBRIC Overall<sup>8</sup> | Sensitivity | 0.633 | 0.717 | 0.133 | 0.832 | 0.795 | 0.847 | 0.790 | 0.799 | 0.865 | 0.866 | 0.602 | 0.599 | 0.771 | 0.776 |
|  | Specificity | 0.884 | 0.994 | 1.000 | 1.000 | 0.999 | 0.994 | 0.999 | 0.995 | 0.995 | 0.995 | 1.000 | 1.000 | 0.999 | 0.999 |
|  | Precision | 0.122 | 0.765 | 0.976 | 0.976 | 0.962 | 0.853 | 0.962 | 0.793 | 0.801 | 0.802 | 0.715 | 0.714 | 0.457 | 0.475 |
|  | Accuracy (MCC) | 0.240 | 0.734 | 0.355 | 0.899 | 0.871 | 0.844 | 0.870 | 0.791 | 0.829 | 0.830 | 0.656 | 0.654 | 0.593 | 0.607 |

\* Dataset time-filtered to eliminate reads from periods of failed skipping, see Supplemental Section S-3.

Figure 2 underscores the importance of such detailed analysis, as simply comparing target- and background-mapping fast5 ratios for odd (10,881:20,761) and even pores (14,312:23,865) can be misleading. Despite an apparent 32% increase in RUBRIC target reads, only 68% of those reads—less than the count of odd target reads— resulted from sequence decisions, while 17% were actively skipped or diverted from the decision process by the threshold filter. The remaining 15% never received a decision, most because they were not reported to RUBRIC by the Event Sampler. We now discuss the read fractions represented in Figure 2, referencing individual results of experiment B2 (Figures 2-3 and Figure 4 (a)) and aggregate results of the other lambda DNA experiments (Table 2, Supplementary Figures S1-S3, S7, and S9-S10).

**Figure 3:**
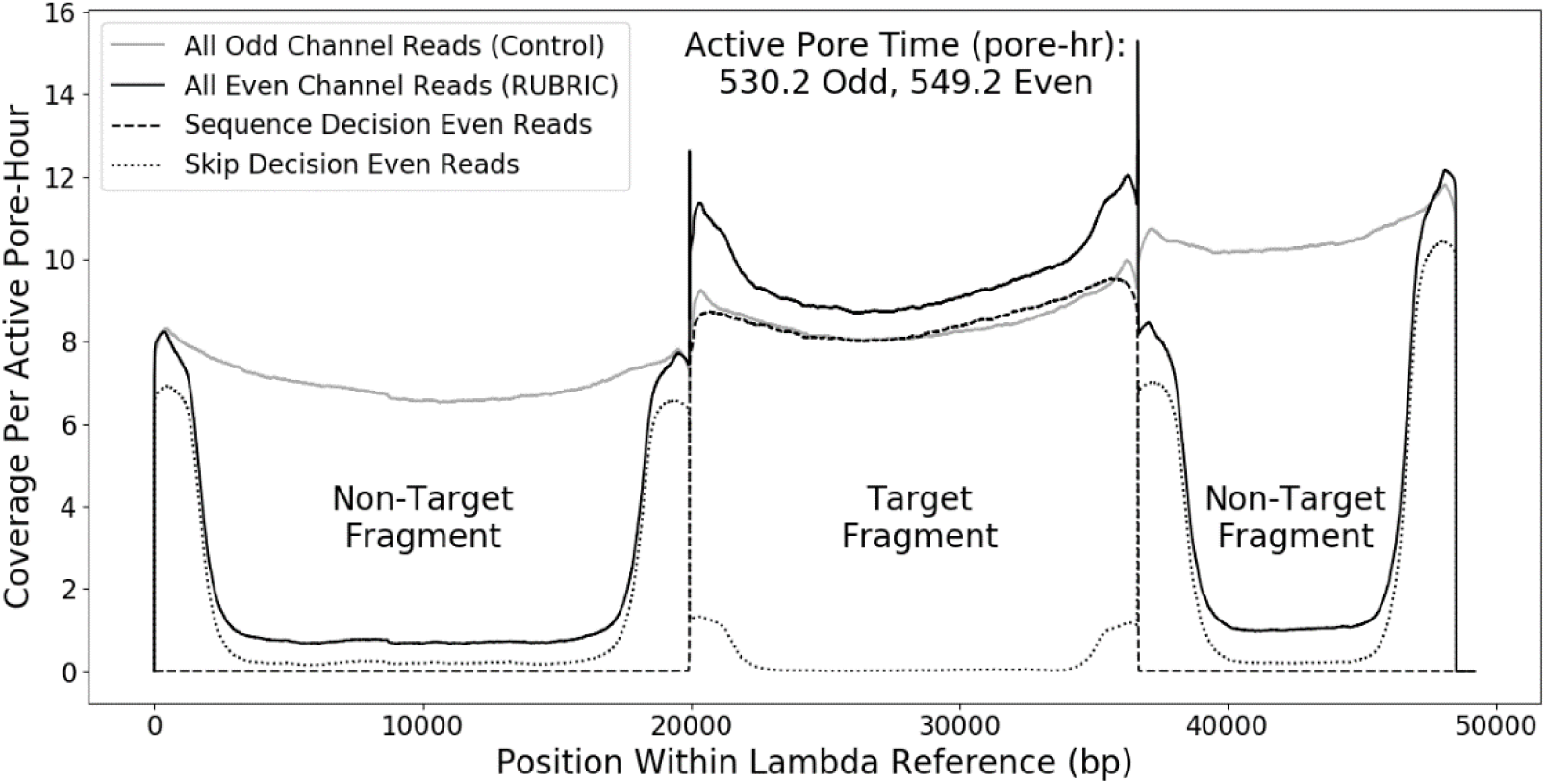
Lambda DNA sequence coverage plot for experiment B2 showing the effect of RUBRIC selection applied to even pore reads in contrast to unselected odd pore reads. Even and odd coverage numbers are normalized by total even and odd active pore times, respectively.

**Figure 4:**
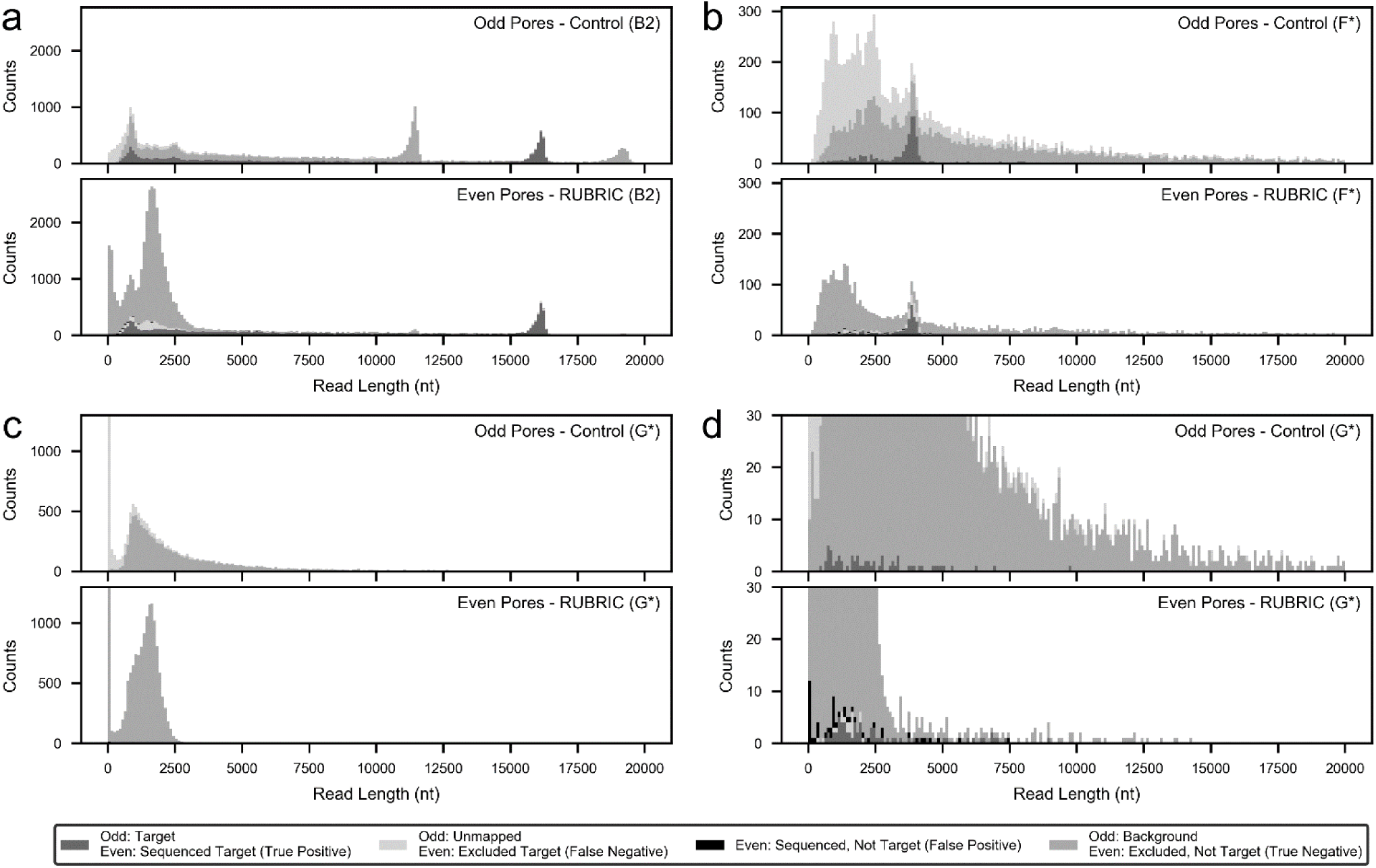
Read length histograms for RUBRIC selection experiments illustrating the distribution of different read types (target, non-target, unmapped) and their fate as a function of RUBRIC selection applied to even numbered pores. Here, reads excluded by the selection process (i.e. not receiving an affirmative sequence decision) include skipped, out-of-threshold, and undecided reads, while reads not mapped to target include those mapped to background/non-target sequence as well as unmappable reads. (a) Lambda DNA experiment B2 showing selection for the middle (nominally ∼17kb) fragment. (b) Example use case dataset F* showing selection for Cas9-excised rDNA from E. coli gDNA. (c, d) Example use case dataset G* showing selection of 1% E. coli gDNA from a background of 99% human gDNA. Supplementary Figure S10 provides more detailed distributions of all read types and categories.

#### Sampled Reads

The character of reads communicated to RUBRIC by the Read Until Event Sampler is best represented by odd pore (control) reads, which exhibited average fragment lengths of 8007 ± 5882 nucleotides (nt) and Albacore quality scores (sequencing_summary.txt-derived “mean_qscore_template”) of 9.52 ± 2.00 for n=214,445 fast5s from N=8 lambda experiments (Supplementary Figure S2).

#### Unsampled Reads

A small percentage (0.62% ± 0.42%, N=8 runs) of reads had fast5 files but lacked Event Sampler entries in the RUBRIC log and were therefore unavailable for selection. These “unsampled” reads typically had quality scores (9.13 ± 2.26, n=34,455 fast5s, N=8 runs) and proportions of target, non-target, and unmappable reads comparable to the sampled control population (Figure 2, Supplementary Figures S2 and S9). The short length (583 ± 206 nt, n=34,455 fast5s, N=8 runs) of most unsampled reads (Supplementary Figure S10), suggests that they may result from DNA transiting the pore within the sampling period of the Event Sampler.

#### Non-Sequence Reads

As in Figure 2, a consistently large proportion of control (odd) sampled reads (89.5% ± 1.89%, N=8 lambda runs) never yielded fast5 sequence files. Pore activity timelines (data not shown) reveal that these “non-sequence” reads typically appear as serial, discretely reported events occurring between identifiable sequence-producing reads. The hypothesis that these non-sequence reads primarily indicate sub-sampling of open pore time (versus degraded DNA, pore fouling, etc.) is reinforced by our observation (data not shown) that setting RUBRIC to unblock all out-of-threshold (predominantly non-sequence) reads produced no apparent change in even pore throughput. A related internal sampling artifact may cause the observed subdivision of long DNA reads^2^.

#### Uncalled Reads

The total number of fast5s that could not be basecalled offline by Albacore was essentially negligible, ranging from 0.0384% (A2) to 0.621% (D) with an average of 0.280% ± 0.246% (N=8 lambda runs) and zero (0) sequence decision fast5s failing to basecall.

#### Mapped & Unmapped Reads

Supplementary Figure S2 shows that odd unmapped reads exhibited significantly lower average quality scores (6.07 ± 1.26, n=35,083 fast5s, N=8 lambda runs) than reads mapping to target or background references (10.21 ± 1.23, n=179,057 fast5s, N=8 lambda runs) and were shorter on average (4082 ± 5556 nt vs. 8789 ± 5625 nt) than corresponding mappable reads.

#### Out-of-Threshold (OOT) Reads

Threshold filter settings (Table 1) were determined empirically from prior run data, requiring updates after any significant sample composition, flowcell batch, library prep, or ONT software changes. Generally, out-of-threshold fast5 quality score averages were about 15% lower than corresponding odd scores (Supplementary Figure S2) and OOT reads about 30% shorter on average. While retrospectively-set thresholds for most mainline experiments successfully excluded 90-97% of ultimately unmappable (especially non-sequence) reads from the decision process, typically diverting >80% of even sampled reads, experiment C showed a lower out-of-threshold proportion (53.7%), rejecting only 56.6% of unmappable reads (Supplementary Figure S9 (f)). This poor threshold selectivity likely accounted for the unusually high in-threshold read/min rate of experiment C (43% higher than B2, Table 2), which in combination with its small queue, may explain its high proportion of undecided reads. Based on C, threshold adjustments in experiment D (Supplementary Figure S9 (g)) produced much improved threshold specificity, precision, and accuracy at the expense of reduced sensitivity (Table 2). Though not optimized when introduced in experiments E1 and E2 (Table 2, Supplementary Figure S9 (h-j)), thresholds based on pore current standard deviation proved superior to those based on mean current because the former helped to mitigate errors associated with current drift and other offsets (Supplementary Section S-2).

#### Undecided & Timeout Reads

The presence of in-threshold reads not receiving skip/sequence decisions typically reflected a computational resource limitation affecting the MinKNOW or RUBRIC PCs. Table 2 indicates the fraction of undecided reads exceeding the 2 second RUBRIC queue timeout period. Excepting outlier experiment C, about 99% of in-threshold reads for mainline lambda experiments received decisions (Table 2). The high in-threshold read rate and poor decision efficiency of experiment C may indicate that as configured the RUBRIC system could effectively process 400-500 decisions/min, beyond which computing resource limitations became significant. Threshold filtering caused undecided reads to differ from control reads mainly in their lower, but variable proportion of non-sequence reads. Because undecided and timeout reads often appeared in localized clusters on the read timeline (see especially Supplementary Figure S5 (d)), this variability may reflect periods of unusually high read throughput that also affected whether fast5s were created by the MinKNOW PC.

#### Sequence Decision Reads

Table 2 details the performance of the RUBRIC decision process in rendering sequence decisions for target mapping reads and skip decisions for non-target reads. For experiment B2, Figure 3 indicates the coverage of lambda (target and non-target) sequence with and without selection, while Figure 4(a) illustrates selection as a function of DNA fragment length. On average for mainline lambda experiments, the decision process correctly excluded 97.7% ± 1.9% (N=5) of non-target reads while capturing 91.4% ± 5.1% (N=5 runs) of available targets, proportions that reflect both basecalling accuracy and the stringency of LAST aligner settings used within the RUBRIC control loop. On average, 98.5% ± 0.6% (N=5) of sequence decision fast5s mapped to target, and even including the typically small proportion of unmapped fast5s (1.5% ± 0.6%), sequence decision quality scores (Supplementary Figures S2-S3) were better on average (10.46 ± 1.36, n=42,191 fast5s) than the control sampled read population (9.51 ± 2.17, n=1,690,891 fast5s). These results suggest that for diagnostic applications, data analysis should focus on sequence decision fast5s and consider other categories (i.e., undecided, unsampled, out-of-threshold, and skipped reads, in that order) only if coverage is lacking.

#### Skip Decision Reads

While skipping ostensibly ejects DNA from the nanopore, on average 46.7% ± 6.1% of mainline experiment skip decisions nevertheless produced fast5s (N=4, excluding outlier C, where the ill-set threshold admitted many non-sequence reads). Skipped-read fast5s occur for two primary reasons. First, when a skip instruction is received, MinKNOW assesses whatever read data has already been acquired and writes it to fast5 if it represents viable sequence (personal communication with ONT staff, 1-9-2018). When skipping is operating correctly with decision times substantially shorter than DNA pore-transit times, this data handling convention produces characteristic truncation of skipped reads visible in the even pore results of Figure 4 and Supplementary Figure S10 as a prominent mound of skipped reads typically centered in the 1500-2500 nt size range. Figure 3 also shows these skip-truncated reads as the higher-coverage “rabbit ear” features (also observed by Loose^33^) at the ends of the non-target lambda fragments. The absence of skip-truncation is an important indication that Read Until DNA rejection is not operating correctly, as discussed in Supplementary Section S-3. Skip decision fast5s may also result when reads transit the pore before a RUBRIC decision can be rendered, whether due to relatively short DNA fragments or long decision times (see Supplementary Section S-6). Unlike skip-truncated reads, which appear only in the even pore results of Figure 4 and the like, reads short enough to escape the decision process in this manner are visible in both odd and even distributions, typically below 1000 nt. In combination, fugitive reads and skip-truncation yielded short average skipped-read lengths of 1373 nt ± 606 nt (n=424,857 fast5s, N=8 lambda runs), while average quality scores were 8.76 ± 2.50 (Supplementary Figure S2).

#### Overall RUBRIC Performance

Table 2 reports absolute target enrichment on both a sequence- and read-basis. Overall, absolute enrichment results were not particularly encouraging, as only mixed sample run G realized both read and sequence enrichment (+15% sequence based on 66 reads for filtered dataset G*, Supplementary Figure S9 (n)), while lambda run B2 showed a nominal gain in read count (2.1%) but slight depletion (1.3%) of target sequence. Other runs saw net reductions in target sequence as great as 24.4% for lambda run E1 and 57.8% for filtered rDNA dataset F* (Supplementary Figure S9 (h) and (l), respectively).

To help understand these results, Supplementary Section S-7 derives a model predicting the likely best-case performance of RUBRIC-style real-time selection for different libraries and computing configurations. In short, because selection only rejects non-target reads, absolute target enrichment is only realized by increasing the total throughput of (even) RUBRIC reads vs. (odd) control reads. Equation 6 in the supplement expresses the maximum absolute enrichment (and throughput enhancement) ratio

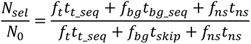

as a function of read fractions (*f*) for target (*t*), background/non-target (*bg*), and non-sequence (*ns*) reads and the characteristic times required to sequence target reads (*t_t_seq_*) and background reads without selection (*t_bg_seq_*), skip background reads with selection (*t_skip_*), and pass non-sequence reads independent of selection (*t_ns_*). As the formula indicates, absolute enrichment is purely a consequence of the time saved by skipping versus sequencing background reads, scaled by their relative prevalence. Furthermore, low pore occupancy (large *f_ns_t_ns_*), as in the experiments described here (Table 2 and Supplementary Table S1), significantly diminishes the benefits of selection. Discrepancies between the empirically observed throughput and absolute enrichment ratios in Table 2 mainly reflect inefficiencies and imperfections in the RUBRIC selection process.

Beyond absolute enrichment, relative enrichment (Table 2) also provides a practical indication of how depleting non-target reads improves the final sequence pool. Computed as the ratio of sequence decision target reads per non-target read divided by the ratio of odd target reads per non-target read, relative enrichment ranges from ∼130x to ∼330x for mainline lambda experiments. This metric underscores the idea that sequence decisions yield such highly purified target-mapping sequence that in most use cases, significant time savings can be realized by analyzing only these reads.

### Example Use Cases

Figure 4 (b) and Supplementary Figures S9 (l) and S10 (l) show the result of RUBRIC selection applied to Cas9-cut *E. coli* gDNA (dataset F*). The target-mapping peak associated with cut rDNA fragments is particularly prominent because 1) *E. coli* has seven copies of the rDNA locus and 2) the AMPure XP beads used in the 1D^2^ library prep provide some positive size selection in the relevant 4-5kb range. While RUBRIC rDNA-mapping reads were reduced 54% versus control, only 3.2% of mappable sequence decision reads mapped to background gDNA versus 89.3% in the control case, yielding relative enrichment of ∼290x. Table 2 reveals suboptimal threshold settings for this run, which realized high specificity but low sensitivity with 38% of the relatively rare target reads falling out-of-threshold. Despite overly aggressive threshold filtering, skip/sequence decisions performed well and had the lowest average decision time (0.23 sec) of any experiment here (Supplementary Figure S7 and Table S1), likely due to the shorter rDNA target reference and low read rates (Table 2) attributable to the relatively dilute library (Table 1).

Figure 4 (c-d) and Supplementary Figures S9 (n) and S10 (o-p) show the result of *E. coli* selection in the mixed human/*E. coli* experiment (dataset G*). Despite LAST-aligning the RUBRIC evaluation window to the entire 4.6 Mb *E. coli* genome for selection, decision times still averaged only 0.91 sec (Supplementary Figure S7 and Table S1). Significantly for this application, aligner stringency was reduced to maximize the number of rare bacterial reads that would be captured, while the evaluation window was doubled to provide additional discrimination between human and bacterial sequence. Consequently, while more sequence decision reads mapped to target (66 vs. 63 control), 42.1% of sequence decision fast5s did not map to target. Moreover, of 84 total even target reads, two were lost to the threshold filter and 17 to skip decisions, as indicated by the comparatively low decision sensitivity, precision, and accuracy for this run. Specificity, however, was comparable to the best seen here, reflecting the comparatively large number of correctly skipped non-target reads. Threshold settings for run G also performed better overall than for any other experiment. Beyond providing nominal absolute target enrichment, the run achieved ∼290x improvement in sequence decision target:non-target ratio due to background depletion of the original 1:99 library.

## Discussion

In this article, we have introduced RUBRIC, a new adaptation of real-time selective sequencing for the Oxford MinION. Unlike the earlier pattern-matching approach^33^, RUBRIC operates in sequence-space, making it possible to leverage the speed, flexibility, and scalability of bioinformatic tools like LAST for selection. Significantly, RUBRIC pre-screening features seek to admit only informative and timely reads to the decision process, reducing computational requirements and enabling real-time basecalling, alignment, and selection of MinION reads without specialized, high-performance computing platforms. While real-time selective sequencing generally provides a means to enrich rare target sequence vs. background without target-specific reagents, primers, or baits, working in sequence-space simplifies the process of choosing, optimizing, and modifying RUBRIC selection targets, all of which can be done on-the-fly based on conventional nucleic acid reference sequences.

We have characterized RUBRIC operation through a series of lambda DNA digest experiments, obtaining limited absolute enrichment of target reads (<2%) but achieving very effective background depletion yielding as much as 330x relative enrichment versus control. The high degree of customization offered by RUBRIC (choice of basecaller/aligner, ratio of RUBRIC to control pores, threshold filter settings, queue size, queue timeout, evaluation window size/offset, aligner settings, etc.) makes it readily adaptable to different sample types, libraries, and computing configurations. Preliminary demonstration experiments have applied RUBRIC to select for CRISPR/Cas9-excised rDNA against a background of *E. coli* gDNA and to select for 1% *E. coli* gDNA against a background of 99% human DNA, achieving absolute target sequence enrichment of 15% in the latter case. To better understand these seemingly modest outcomes, we have proposed a model estimating the likely upper bounds on real-time selection performance and have found our results to be largely consistent with its predictions. This analysis suggests that the limited target enrichment we have seen to date is less a consequence of the speed or fidelity of our method than the relatively high rate of MinION pore vacancy, which critically limits the gains that can be realized by real-time selection.

Future work will focus on optimizing RUBRIC performance and applying the method to clinically and diagnostically relevant sample types (e.g., host/pathogen mixtures), where selection can provide the greatest benefits. In such applications, accumulating RUBRIC sequence decision reads could itself provide a rapid, presumptive diagnostic result, given sufficient specificity. These reads could also be used to prioritize which fast5s should receive concurrent full strand basecalling and analysis during sequencing, potentially shortening time to identification. With these goals in mind, we will seek to improve our library preparations to increase pore occupancy and DNA fragment length, both of which should substantially improve RUBRIC performance fragment length, both of which should substantially improve RUBRIC performance based on our model predictions. To avoid the pitfalls of retrospectively setting the RUBRIC threshold filter, we plan to automate this process, perhaps using real-time RUBRIC decision and mapping results to iteratively adjust the filter throughout each run. We also expect to migrate RUBRIC to the latest release of the Read Until developer API (v2), adapt the method for raw data or GPU basecalling (e.g., with ONT’s Scrappie or Guppy callers, respectively), and explore its application to MinION direct RNA sequencing.

## Acknowledgements

Thanks to Matt Loose of the University of Nottingham for his early guidance in scaling the Read Until learning curve and to Julian Atienza, George Pimm, Richard Ronan, and Chris Wright of ONT for their advice and support in providing access to pre-release versions of the Read Until API. Thanks also to Steve Branda and Joe Schoeniger for their work in establishing and advising this project. This work was supported by the Laboratory Directed Research and Development (LDRD) program at Sandia National Laboratories, a multi-mission laboratory managed and operated by National Technology and Engineering Solutions of Sandia, LLC, a wholly owned subsidiary of Honeywell International, Inc., for the U.S. Department of Energy’s National Nuclear Security Administration under contract DE-NA0003525.

## Author Contributions

H.S.E. wrote the code, ran sequencing, performed analysis, and contributed to the manuscript. R.K. conceived and designed the project, contributed to the code, performed analysis, and contributed to the manuscript. A.S. performed sample extractions, digests, and library preparations, ran sequencing, and contributed to the manuscript. S.W.B. performed CRISPR/Cas9 guide design and developed digests. K.D.P. conceived and designed the project and contributed to the manuscript. M.S.B. conceived, designed, and led the project, performed analysis, developed the model, and wrote the manuscript.

## Additional Information

### Materials & Correspondence

Please address correspondence and inquiries to M.S.B.

### Code Availability

The RUBRIC code and Nanonet basecaller are publicly available at https://github.com/harrisonedwards/RUBRIC. Please check the associated Github Issues page and post any problems or questions there before contacting the authors directly. The Read Until v1 API needed to run RUBRIC can be obtained directly from ONT through their Developer License Agreement.

### Data Availability

RUBRIC logfiles and associated metadata are publicly available at https://doi.org/10.25739/71ne-xy91, while all MinION-produced fast5 sequence files are available at https://www.ncbi.nlm.nih.gov/bioproject/PRJNA491460.

## References

1 Jain, M., Olsen, H. E., Paten, B. & Akeson, M. The Oxford Nanopore MinION: delivery of nanopore sequencing to the genomics community. Genome Biology 17, doi:10.1186/s13059-016-1103-0 (2016).

2 Payne, A., Holmes, N., Rakyan, V. & Loose, M. Whale watching with BulkVis: A graphical viewer for Oxford Nanopore bulk fast5 files. bioRxiv, doi:10.1101/312256 (2018).

3 Garalde, D. R. et al. Highly parallel direct RNA sequencing on an array of nanopores. Nat. Methods 15, 201–206, doi:10.1038/nmeth.4577 (2018).

4 Cornelis, S., Gansemans, Y., Deleye, L., Deforce, D. & Van Nieuwerburgh, F. Forensic SNP genotyping using nanopore MinION sequencing. Sci Rep 7, doi:10.1038/srep41759 (2017).

5 Tarumoto, N. et al. Use of the Oxford Nanopore MinION sequencer for MLST genotyping of vancomycin-resistant enterococci. Journal of Hospital Infection 96, 296–298, doi:10.1016/j.jhin.2017.02.020 (2017).

6 Giolai, M. et al. Comparative analysis of targeted long read sequencing approaches for characterization of a plant’s immune receptor repertoire. BMC Genomics 18, doi:10.1186/s12864-017-3936-7 (2017).

7 Hyeon, J.-Y. et al. Quasimetagenomics-based and real-time-sequencing-aided detection and subtyping of Salmonella enterica from food samples. Applied and Environmental Microbiology 84, doi:10.1128/aem.02340-17 (2018).

8 Brown, B. L., Watson, M., Minot, S. S., Rivera, M. C. & Franklin, R. B. MinION (TM) nanopore sequencing of environmental metagenomes: a synthetic approach. Gigascience 6, doi:10.1093/gigascience/gix007 (2017).

9 Goordial, J. et al. In situ field sequencing and life detection in remote (79 degrees 26’ N) Canadian high arctic permafrost ice wedge microbial communities. Frontiers in Microbiology 8, doi:10.3389/fmicb.2017.02594 (2017).

10 Norris, A. L., Workman, R. E., Fan, Y. F., Eshleman, J. R. & Timp, W. Nanopore sequencing detects structural variants in cancer. Cancer Biology & Therapy 17, 246–253, doi:10.1080/15384047.2016.1139236 (2016).

11 Suzuki, A. et al. Sequencing and phasing cancer mutations in lung cancers using a long-read portable sequencer. DNA Res. 24, 585–596, doi:10.1093/dnares/dsx027 (2017).

12 Ashton, P. M. et al. MinION nanopore sequencing identifies the position and structure of a bacterial antibiotic resistance island. Nature Biotechnology 33, 296-+, doi:10.1038/nbt.3103 (2015).

13 Schmidt, K. et al. Identification of bacterial pathogens and antimicrobial resistance directly from clinical urines by nanopore-based metagenomic sequencing. Journal of Antimicrobial Chemotherapy 72, 104–114, doi:10.1093/jac/dkw397 (2017).

14 Jain, M. et al. Nanopore sequencing and assembly of a human genome with ultra-long reads. Nature Biotechnology 36, 338-+, doi:10.1038/nbt.4060 (2018).

15 Michael, T. P. et al. High contiguity Arabidopsis thaliana genome assembly with a single nanopore flow cell. Nat. Commun. 9, 8, doi:10.1038/s41467-018-03016-2 (2018).

16 Tyson, J. R. et al. MinION-based long-read sequencing and assembly extends the Caenorhabditis elegans reference genome. Genome Research 28, 266–274, doi:10.1101/gr.221184.117 (2018).

17 Minh Duc, C. et al. Streaming algorithms for identification of pathogens and antibiotic resistance potential from real-time MinION (TM) sequencing. Gigascience 5, doi:10.1186/s13742-016-0137-2 (2016).

18 Hoenen, T. et al. Nanopore sequencing as a rapidly deployable Ebola outbreak tool. Emerg. Infect. Dis 22, 331–334, doi:10.3201/eid2202.151796 (2016).

19 Quick, J. et al. Real-time, portable genome sequencing for Ebola surveillance. Nature 530, 228–232, doi:10.1038/nature16996 (2016).

20 Quick, J. et al. Multiplex PCR method for MinION and Illumina sequencing of Zika and other virus genomes directly from clinical samples. Nature Protocols 12, 1261–1276, doi:10.1038/nprot.2017.066 (2017).

21 Votintseva, A. A. et al. Same-day diagnostic and surveillance data for tuberculosis via whole-genome sequencing of direct respiratory samples. Journal of Clinical Microbiology 55, 1285–1298, doi:10.1128/jcm.02483-16 (2017).

22 Imai, K. et al. An innovative diagnostic technology for the codon mutation C580Y in kelch13 of Plasmodium falciparum with MinION nanopore sequencer. Malar. J. 17, 11, doi:10.1186/s12936-018-2362-x (2018).

23 Russell, J. A. et al. Unbiased strain-typing of arbovirus directly from mosquitoes using nanopore sequencing: a field-forward biosurveillance protocol. Sci Rep 8, 12, doi:10.1038/s41598-018-23641-7 (2018).

24 Greninger, A. L. et al. Rapid metagenomic identification of viral pathogens in clinical samples by real-time nanopore sequencing analysis. Genome Medicine 7, doi:10.1186/s13073-015-0220-9 (2015).

25 Quick, J. et al. Rapid draft sequencing and real-time nanopore sequencing in a hospital outbreak of Salmonella. Genome Biology 16, doi:10.1186/s13059-015-0677-2 (2015).

26 Hagemann, I. S., Cottrell, C. E. & Lockwood, C. M. Design of targeted, capture-based, next generation sequencing tests for precision cancer therapy. Cancer Genetics 206, 420–431, doi:10.1016/j.cancergen.2013.11.003 (2013).

27 Forbes, J. D., Knox, N. C., Ronholm, J., Pagotto, F. & Reimer, A. Metagenomics: the next culture-independent game changer. Frontiers in Microbiology 8, 21, doi:10.3389/fmicb.2017.01069 (2017).

28 Taylor-Brown, A., Madden, D. & Polkinghorne, A. Culture-independent approaches to chlamydial genomics. Microbial genomics, doi:10.1099/mgen.0.000145 (2018).

29 Brinkmann, A. et al. Development and preliminary evaluation of a multiplexed amplification and next generation sequencing method for viral hemorrhagic fever diagnostics. Plos Neglect. Trop. Dis. 11, doi:10.1371/journal.pntd.0006075 (2017).

30 Karamitros, T. & Magiorkinis, G. A novel method for the multiplexed target enrichment of MinION next generation sequencing libraries using PCR-generated baits. Nucleic Acids Research 43, 11, doi:10.1093/nar/gkv773 (2015).

31 Kumar, A., Murthy, S. & Kapoor, A. Evolution of selective-sequencing approaches for virus discovery and virome analysis. Virus Research 239, 172–179, doi:10.1016/j.virusres.2017.06.005 (2017).

32 Shin, G. et al. CRISPR-Cas9-targeted fragmentation and selective sequencing enable massively parallel microsatellite analysis. Nat. Commun. 8, doi:10.1038/ncomms14291 (2017).

33 Loose, M., Malla, S. & Stout, M. Real-time selective sequencing using nanopore technology. Nat. Methods 13, 751–754, doi:10.1038/nmeth.3930 (2016).

34 Yamamoto, T. & Fujii, T. Nanofluidic single-molecule sorting of DNA: a new concept in separation and analysis of biomolecules towards ultimate level performance. Nanotechnology 21, doi:10.1088/0957-4484/21/39/395502 (2010).

35 Cipriany, B. R. et al. Real-time analysis and selection of methylated DNA by fluorescence-activated single molecule sorting in a nanofluidic channel. Proceedings of the National Academy of Sciences of the United States of America 109, 8477–8482, doi:10.1073/pnas.1117549109 (2012).

36 Bain, F. E., Wu, C. G. & Spies, M. Single-molecule sorting of DNA helicases. Methods 108, 14–23, doi:10.1016/j.ymeth.2016.05.009 (2016).

37 Masutani, B. & Morishita, S. A framework and an algorithm to detect low-abundance DNA by a handy sequencer and a palm-sized computer. Bioinformatics, doi:10.1093/bioinformatics/bty663 (2018).

38 Kielbasa, S. M., Wan, R., Sato, K., Horton, P. & Frith, M. C. Adaptive seeds tame genomic sequence comparison. Genome Research 21, 487–493, doi:10.1101/gr.113985.110 (2011).

39 Krishnakumar, R. et al. Systematic and stochastic influences on the performance of the MinION nanopore sequencer across a range of nucleotide bias. Sci Rep 8, 13, doi:10.1038/s41598-018-21484-w (2018).

40 Anders, C. & Jinek, M. In Vitro Enzymology of Cas9. Methods in Enzymology 546, 1–20, doi:10.1016/b978-0-12-801185-0.00001-5 (2014).

