## Supplementary Information for "Real-Time Selective Sequencing with RUBRIC: Read Until with Basecall and Reference-Informed Criteria"

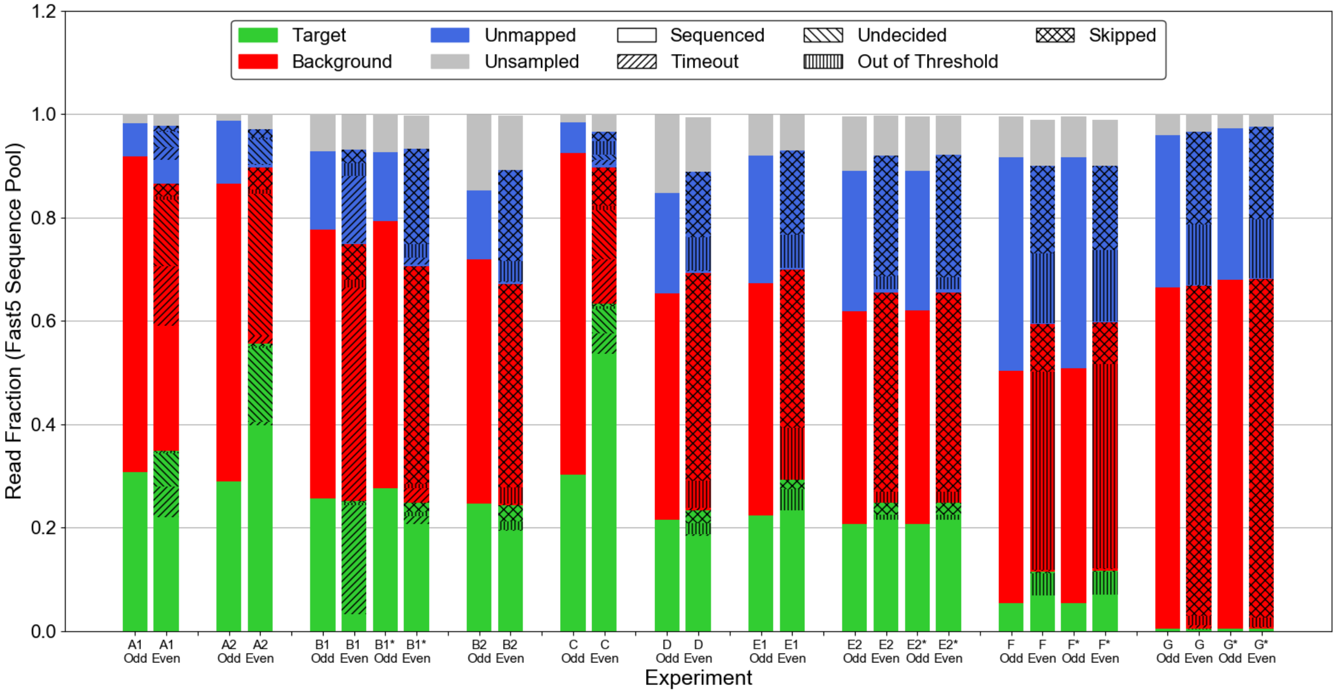


*Figure S1: Compilation of fast5 sequence results for all experiments and time-filtered datasets (*). Shading indicates sampled read mapping and unsampled reads, while fill pattern for sampled reads indicates whether reads received a skip or sequence decision or received no decision due to the threshold filter, queue timeout, or other factors.*

### S-1. Comparative Experimental Metrics

Referencing Table 2 and our Sankey data flow analysis (Figure 2, Supplementary Figure S9), we assess the performance of the threshold filter, the decision process, and the overall end-to-end RUBRIC selection process in terms of binary classifiers, yielding true positive (TP), true negative (TN), false positive (FP), and false negative (FN) counts for each element of the RUBRIC workflow. For purposes of the threshold filter, which primarily serves to exclude unmappable and non-sequence reads from the decision process while admitting mappable reads, true positives are counted as in-threshold mappable reads (both target and background), true negatives as out-of-threshold unmappable reads (including non-sequence and uncalled), false positives as in-threshold unmappable reads, and false negatives as out-of-threshold mappable reads. Within the decision process, which accepts target mapping reads and rejects the rest, true positives are counted as sequence decision target-mapping reads, true negatives as skip decision reads not mapping to target (including non-sequence, uncalled, unmapped, and background-mapping reads), false positives as sequence decision reads not mapping to target, and false negatives as skip decision reads mapping to target. The overall RUBRIC process also discriminates target from non-target reads, and consequently its performance metrics consider true positives as sequence decision reads mapping to target, true negatives as all even pore sampled non-sequence-decision reads not mapping to target (including out-of-threshold, undecided, and skipped reads), false positives as sequence decision reads not mapping to target, and false negatives as all even pore sampled non-sequence-decision reads mapped to target.

RUBRIC Performance metrics given in Table 2 are defined in the conventional manner for binary classification, where for each portion of the workflow noted above sensitivity = TP/(TP+FN), selectivity = TN/(TN+FP), and precision = TP/(TP+FP). Accuracy expressed as (TP+TN)/(TP+TN+FP+FN) proves to be potentially misleading, particularly in experiments with dramatically different positive and negative counts. Accordingly, we compare overall accuracies instead with the Matthews Correlation Coefficient (MCC) given by:

$$MCC = \frac{(TP \times TN) - (FP \times FN)}{\sqrt{(TP+FP)(TP+FN)(TN+FP)(TN+FN)}}$$

Which ranges between -1 (completely inaccurate selection) and 1 (perfect selection). Note that all binary classifier performance metrics in Table 2 implicitly assume that skip decisions do not affect whether reads become fast5 files, potentially exaggerating the number of false positive threshold results, true negative decision results, and true negative RUBRIC results to an unknown degree if this assumption proves to be incorrect.

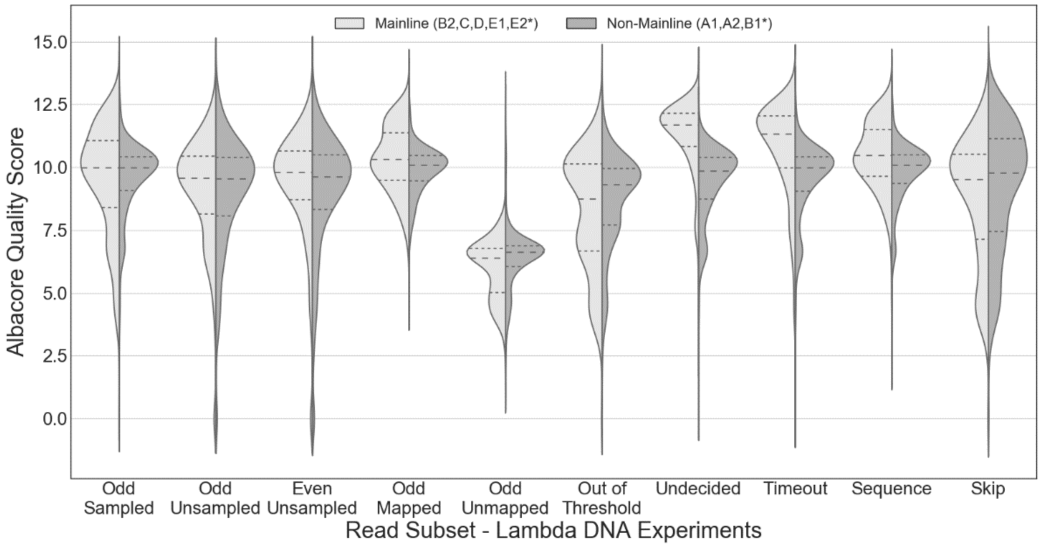


*Figure S2: Quality score distributions for different read subsets of the pooled set of mainline lambda DNA experiments (N=5) in comparison to the set of preliminary, non-mainline experiments (N=3). Dashed lines indicate the quartiles within each distribution.*

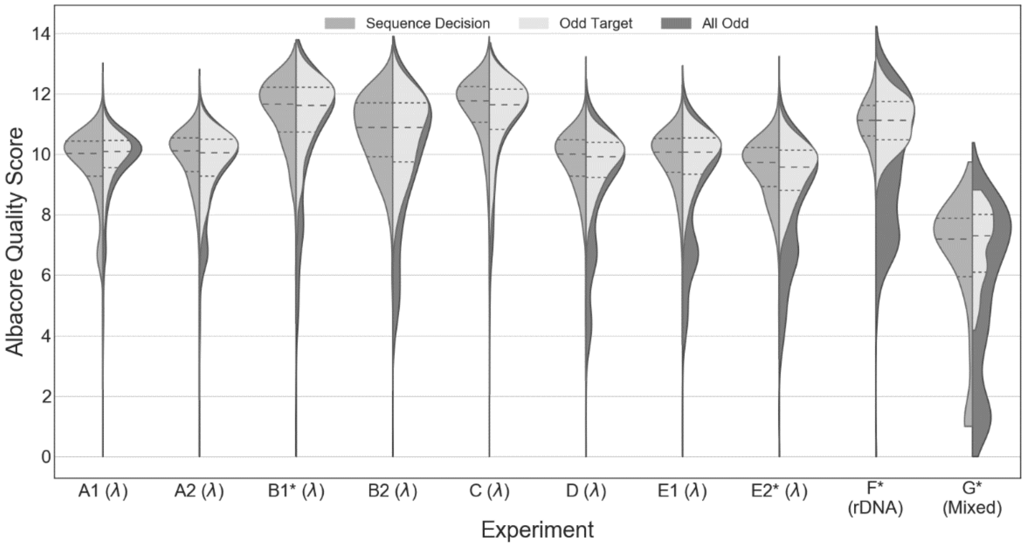


*Figure S3: Overall quality score distributions for even pore sequence decision reads and odd pore (control) reads by experiment for lambda DNA, Cas9-excised E. coli rDNA, and mixed human/E. coli gDNA selection experiments. Datasets B1*, E2*, F*, and G* are filtered to remove periods of failed skipping as described in Section S-3, though the resulting difference to these distributions is minimal. Run G was basecalled with a later version of Albacore, so its scores may not be directly comparable to the other runs.*

### S-2. Threshold Filter Settings

Corresponding to mixed sample dataset G*, Supplementary Figure S4 provides the best available illustration of threshold filter operation (see also Supplementary Figure S9 (n)), highlighting in particular the tradeoffs between admitting as many mappable reads as possible while excluding non-sequence, uncallable, and unmappable reads. Pore current metrics were only logged for out-of-threshold reads prior to experiment G. As noted in the main article, threshold filter settings for a given run were determined retrospectively based on prior datasets, which is why they were not always well-optimized. Initial experiments utilized a threshold based on calculating the mean pore current for the RUBRIC evaluation window, but in later iterations a regression model showed that using standard deviation alone—as opposed to mean alone or a combination of mean and standard deviation—was the best predictor of whether a read would ultimately be mappable (data not shown). In addition to compensating for drift and flowcell-to-flowcell or run-to-run variations in pore current, the use of the standard deviation-based (rather than mean current-based) threshold should also help minimize the effect of programmed 5 mV sequencer voltage adjustments[2] on threshold efficiency over the course of longer runs.

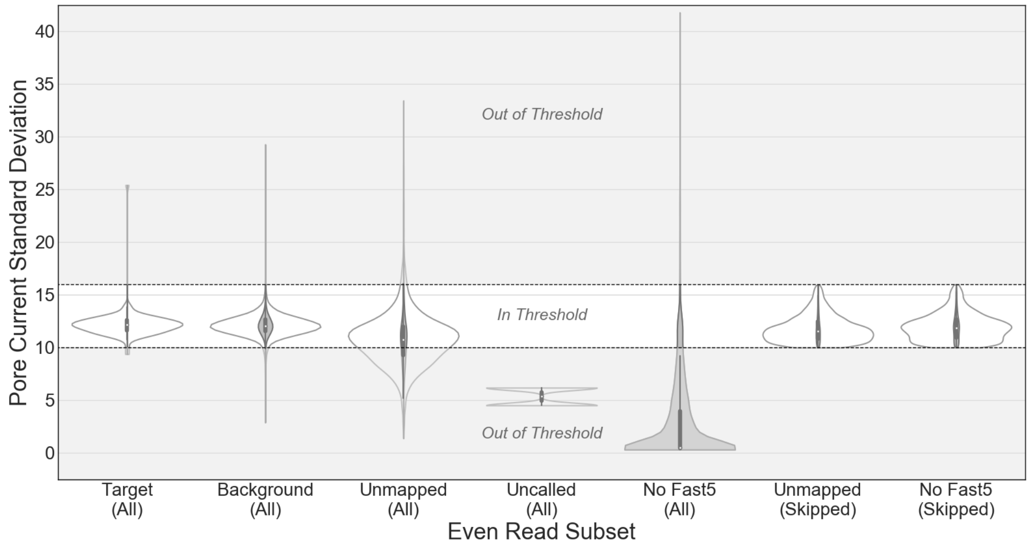


*Figure S4: Distribution of measured even pore current standard deviations (in picoamps) for all mappable and unmappable fast5s as well as non-sequence (no fast5) reads for mixed dataset G* (skip-fail filtered). Dotted lines indicate the position of upper and lower threshold limits, with reads falling above or below this band classified as out-of-threshold. White outer violin profiles show the shape of the distribution independent of read count (fixed width), while inner gray profiles indicate relative counts across the seven categories for the run.*

### S-3. Read Until Event Sampler and Skip Failures

Two significant problems were observed in the course of working with the beta version of the Read Until API and Event Sampler. First, in many experiments the Event Sampler would simply stop communicating with RUBRIC for no apparent reason, sometimes hours before the end of the sequencing run, as reflected in Table 1 by the mismatch between MinION run and Event Sampler times for some experiments. Accordingly, all results discussed in this article reflect only data collected while the Event Sampler was communicating with RUBRIC, though complete datasets are available as noted in the Methods.

The second significant pathology observed in some RUBRIC experiments was the apparent intermittent failure of the DNA rejection / pore unblocking process to respond appropriately to RUBRIC skip decisions. While these skip-failures affected some runs in their entirety (data not shown), closer examination of other runs revealed discrete and readily identifiable time intervals during which the Read Until skipping mechanism had simply failed to reject any DNA, as shown in Supplementary Figure S5 for experiments B1, E2, F, and G. Because these periods of failed skipping potentially introduce uncharacteristic anomalies to our analyses, we distinguish full run data (B1, E2, F, and G) from datasets B1*, E2*, F*, and G* that have been time-filtered to eliminate all reads from the intervals during which skipping did not result in read rejection/truncation. Unless otherwise noted, aggregate results generally refer to these filtered datasets. Differences between filtered and unfiltered datasets are indicated in Tables 1 and 2, Supplementary Table S1, and Supplementary Figures S1, S9, and S10.

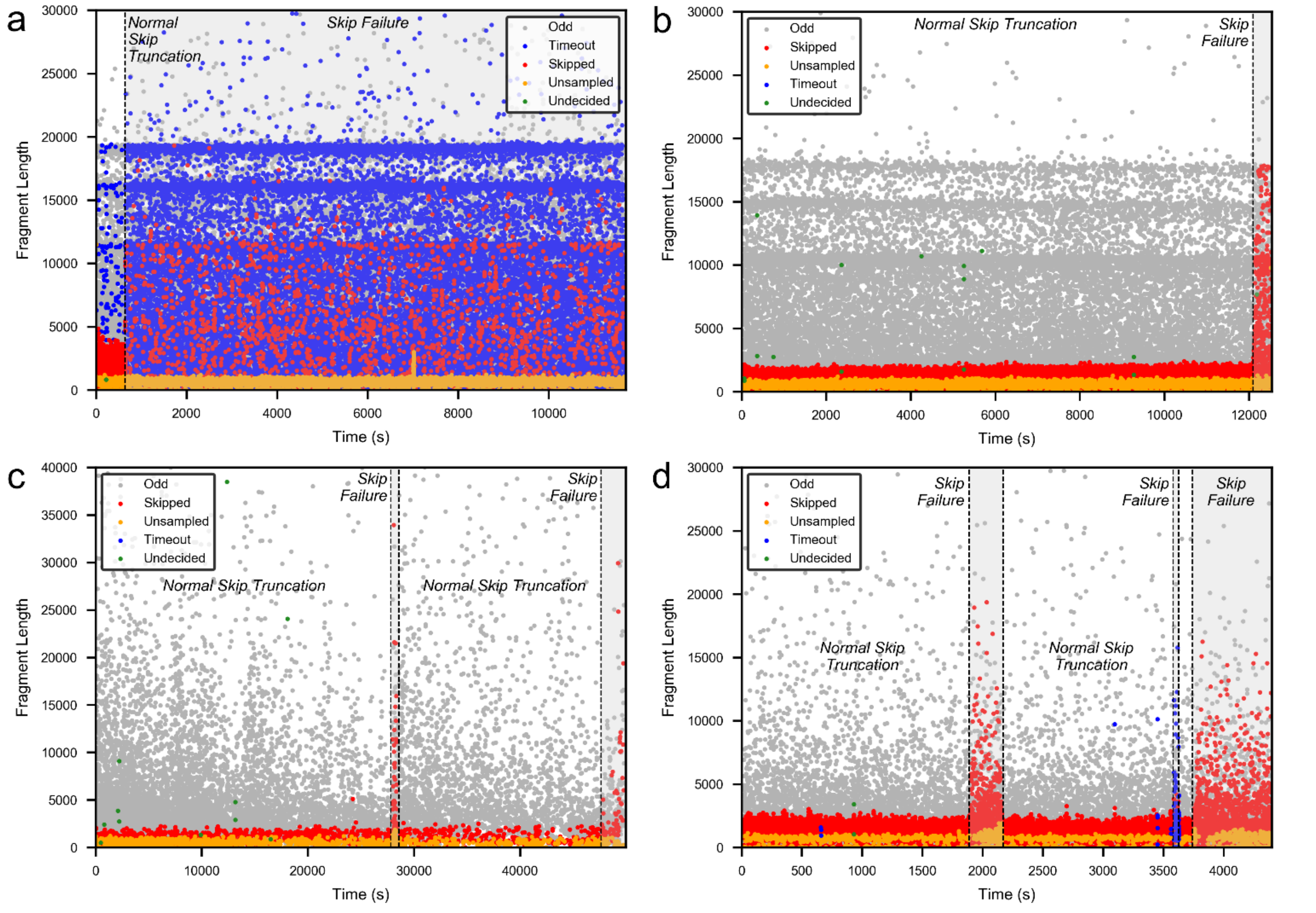


*Figure S5: Scatter plots of read lengths over time for experiments (a) B1, (b) E2, (c) F, and (d) G illustrating the periods of noticeable skip failure indicated by the sudden increases in skipped read lengths. Periods of normal skip truncation in each case form the basis for time-filtered datasets B1*, E2*, F*, and G*.*

In the case of pre-Safe Mode run B1 (Supplementary Figure S5 (a)), only the first 11 minutes of the run exhibited effective skipping, after which a catastrophic resource limitation or communication failure caused the vast majority of even pore reads to time-out of the decision process while decision times spiked as high as 40 seconds. Run E2 (Supplementary Figure S5 (b)) saw a skipping failure around the 200 minute mark that appeared to be uncorrelated to any significant change in either decision times or the incidence of undecided, timeout, or unsampled reads, though the Event Sampler failed altogether six minutes later. Run F (Supplementary Figure S5 (c)) showed two short intervals lacking skip-truncation around the 7.7 hr and 13.3 hr marks, the first lasting about 5 minutes and the second preceding final Event Sampler failure about 30 minutes later, but neither appeared to be associated with unusual decision times or undecided/unsampled reads. Lastly, experiment G (Supplementary Figure S5 (d)) exhibited perhaps the most complex behavior related to skip failures. In this run, three distinctly recognizable intervals of non-truncated skipped reads are present, commencing at the 31 min mark (~5 min duration), the 60 min mark (~1 min duration), and the 63 min mark, with the last interval again immediately preceding the failure of the Event Sampler 11 minutes later. The first and last periods of skipping failure are similar and both encompass a brief introductory period of decision times skewing shorter with reduced unsampled read counts followed by a return to the typical decision time distributions and increased unsampled read density. The second, shortest, period without skipping correlates to both much longer decision times (exceeding 8 s) and a high incidence of reads that time-out of the RUBRIC decision process. While some of these failure modes may be precipitated by reaching the computing resource limits of either the MinKNOW laptop or RUBRIC desktop, others appear to be attributable to errors or instabilities in the beta implementation of the Read Until API (v1) and Event Sampler themselves.

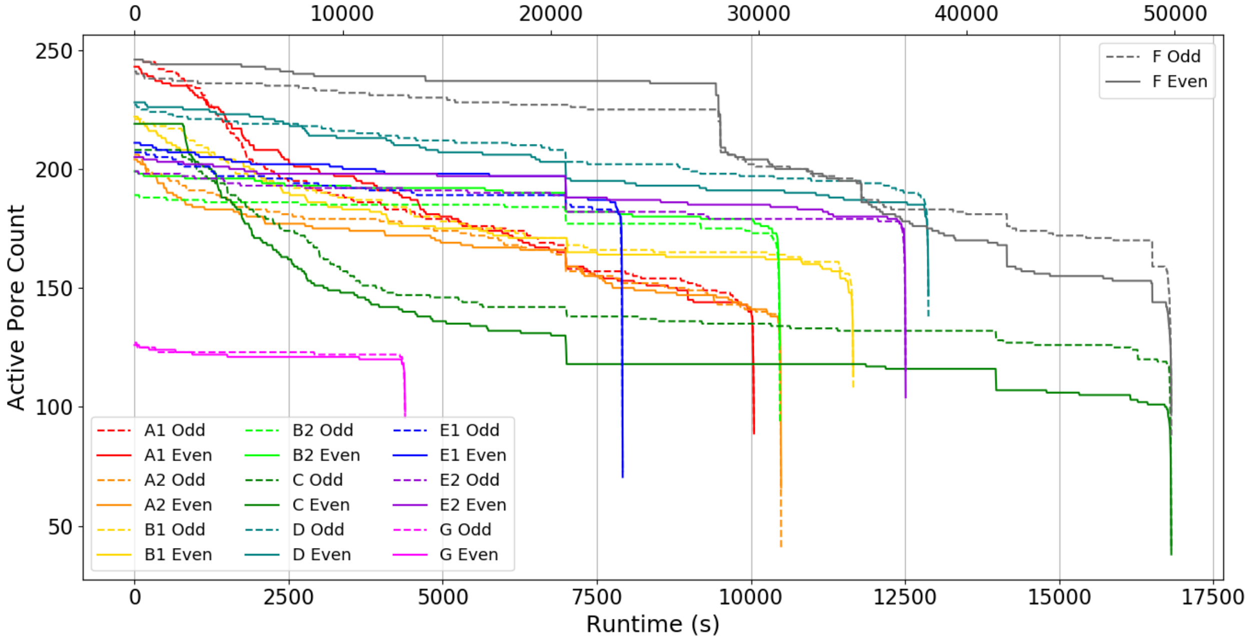


*Figure S6: Pore lifetime with and without RUBRIC selection. Odd (control) pore counts over time reflect typical pore attrition while even pores illustrate the effect of RUBRIC-controlled active unblocking on pore viability. Long experiment F is plotted against the upper x-axis, while all other experiments are plotted against the lower axis.*

### S-4. Pore Lifetime

One significant question regarding the viability of RUBRIC-style real-time selection is whether the act of repeatedly reversing pore polarity (unblocking) to reject non-target DNA degrades pore performance over time or leads to accelerated pore attrition. Supplementary Figure S6 addresses this question, showing even and odd active pore counts for each experiment as a function of time. While nearly all runs end with fewer even pores than odd pores, the rate of pore attrition (slope of the line) is remarkably similar for both skipping and non-skipping pores. Interestingly, large step reductions in even pore counts occur every 2 hours, perhaps coinciding with the 2-hour, 5 mV voltage adjustment interval of the sequencer[2] or the multiplex (mux) group changeover, although it was our understanding that the latter occurs on a significantly longer interval (~24 hours). If a mux changeover is to blame, this may indicate that the act of unblocking the active pore in each bank of four degrades the other inactive pores in that set, a fact which is only revealed when the next group of pores is activated. Another interesting observation: the period of skip failure visible near the 28,000 sec mark in Supplementary Figure S5 (c) appears to coincide with the large decrease in experiment F active pore count shown in Supplementary Figure S6. It is also worth noting that in our experience with the current generation of MinION flowcells, pore attrition—and not library depletion—appears to be the main factor contributing to reduced read counts over time, at least for runs less than 24 hr in duration.

### S-5. Frozen Libraries

As noted above, the library for experiment C was made at the same time as the B1/B2 library and frozen at -20 °C for 1 day before use. Similarly, E1/E2 was prepared with D and frozen for 2 days before use. Supplementary Figure S3 shows no significant differences in the odd read quality score distributions of these two fresh/frozen pairs. As Table 2 shows, the frozen libraries do appear to yield substantially fewer odd reads per pore per minute, a pattern that also persists on a likely more relevant read/pore-min/(ng of input DNA) basis, showing a 45% decrease between runs B2 and C and a 53% decrease for E1 versus run D. Despite these differences in read rate/yield, comparing Supplementary Figure S10 (e) to (f) and Supplementary Figure S10 (g) to (h) shows no evidence of significant DNA fragmentation or read length bias obviously attributable to freezing. In fact, the run C size distribution shows a greater proportion of longer reads than B2. These results suggest that preparing and freezing libraries for later sequencing can be done without dramatically altering library quality or content, though sequence coverage and overall throughput may suffer, perhaps due to damage/degradation of the DNA-tether-motor complex.

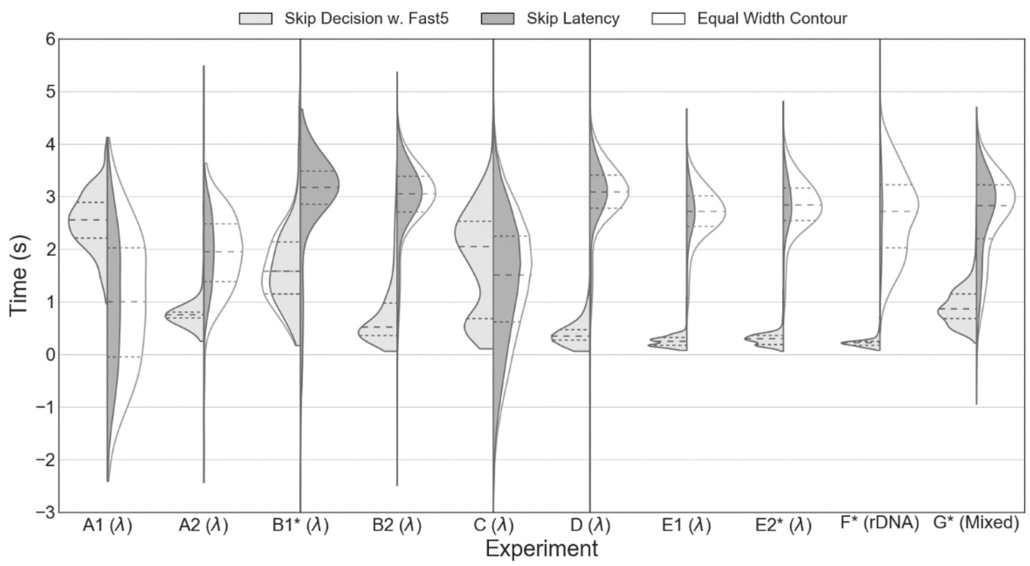


*Figure S7: RUBRIC skip decision time and skip decision latency distributions for real-time selective sequencing experiments. Skip latency is calculated as the difference between the duration of a skipped read and its RUBRIC decision time, corresponding to the sum of any delays in the read skipping timeline that occur either before or after the RUBRIC process itself. Negative "latency" numbers indicate reads that likely received skip decisions after they had already departed the nanopore.*

### S-6. RUBRIC Decision Times & Read Until Latency

Supplementary Figure S7 shows the RUBRIC decision time distributions and calculated skip decision latency times for each experiment described in the article. Decision times are primarily dependent on computing resource availability, the size of the evaluation window that is to be basecalled in real-time, and the size of the target reference sequence used for LAST alignment. Because all reads receiving a decision go through the same basecall and alignment process, there is no difference between skip and sequence decision times. Skip latency, as shown in the figure, is the difference between the Albacore-reported read duration for a given skip decision read and its corresponding RUBRIC decision time. This latency likely represents the effect of 1) any delay between DNA docking at a pore and the Event Sampler reporting that event to RUBRIC and/or 2) any delay between RUBRIC communicating a skip decision and MinKNOW reversing pore polarity. Reads with a duration shorter than the RUBRIC skip decision time (appearing as negative "latencies" on the chart) most likely received decisions after they had already departed the nanopore. Depending on the balance between fragment size and decision time, shortening the RUBRIC queue timeout period may reduce the incidence of such *post hoc* decisions, conserving decision process computing resources at the expense of increasing the number of undecided (timeout) reads. As Supplementary Figure S7 indicates, with the exception of run C, which had a number of unusually long decision times, skip latency times were both substantially longer than RUBRIC decision times and fairly narrowly distributed around a mean of about 3 sec. Unclear is how these latency times relate to the observed 2 second duration of pore unblocking[2]. For a sequencer that is nominally operating at a DNA translocation rate of 450 bases/s, these internal latencies impose a lower limit on the size of DNA fragments that can be effectively selected, regardless of RUBRIC decision time performance or additional optimization.

### S-7. Estimating the Limits of Real-Time Selective Sequencing Performance

Based on our work characterizing the performance of the RUBRIC method, we now propose a simple model providing a means to estimate the generalized performance limits of real-time selection processes like RUBRIC. Central to our analysis is the recognition that on average, the amount of time that a sequencing DNA strand occupies a given nanopore is equal to its length divided by the translocation rate, nominally 450 bases/s with current technology. Accordingly, we posit that the primary benefit of selective sequencing is that it allows some read types to be processed more quickly (i.e., when skipped) than they would be in the absence of selection.

For simplicity, we address three primary read types encountered by the nanopore: 1) reads corresponding to target sequence that are to be sequenced, 2) non-target or background-mapping reads that are to be skipped, and 3) non-sequence or open pore reads that, despite being registered by the Event Sampler, never translate into fast5 sequence files. As noted in the main article, non-sequence reads are believed to reflect the amount of time that a pore is unoccupied, sampled at somewhat regular intervals. Based on our observations, these non-sequence reads can account for 85-99% of the total reported read population and 65-98% of all active pore time (Table 2, Supplementary Table S1), with the remainder of active pore time split between target and non-target reads in essentially the same ratio that their sequences exist in the library. It should be noted that while unmapped reads likely originate as either low quality target or background sequence, they are counted as non-target reads for modeling and performance assessment purposes.

Within a given run, the observed proportions of read types in the read pool reflect the underlying probabilities that an unoccupied nanopore will encounter a particular kind of read, probabilities that themselves reflect the composition of the library (target:background ratio, DNA concentration) and are therefore constant and independent of any selection method. With these assumptions in mind, considering first the case of selective sequencing (*sel*), the total number of target (*t*), background (*bg*), and non-sequence (*ns*) reads obtained in an experiment is given by

$$N_{sel} = n_{t\_sel} + n_{bg\_sel} + n_{ns\_sel} \quad (1)$$

Dividing by $N_{sel}$ we express each read category as its fraction (*f*) of the total read population upstream of the decision process

$$1 = \frac{n_{t\_sel}}{N_{sel}} + \frac{n_{bg\_sel}}{N_{sel}} + \frac{n_{ns\_sel}}{N_{sel}} = f_t + f_{bg} + f_{ns} \quad (2)$$

Importantly, we reiterate that while fast5 counts and proportions downstream may be affected by selection in ways that are hard to predict *a priori*, selection has no effect on these purely library-dependent upstream read fractions. In the bounding case of perfect selection, all target reads will be sequenced and all non-target reads skipped, each requiring on average $t_{t\_seq}$ and $t_{skip}$ seconds of pore-time, respectively, per read. Generally, $t_{t\_seq}$ approximates the average target fragment length divided by the average pore translocation rate (e.g., 450 bases/s), while $t_{skip}$ reflects the total time required for the selection method to assess and implement the skip decision (e.g., RUBRIC decision time plus MinKNOW latencies). We likewise define a characteristic average time $t_{ns}$ associated with non-sequence reads, which are reported by the Event Sampler as discrete events even though they frequently represent subdivisions of continuous intervals of pore vacancy. The cumulative pore-time (*T*) needed to obtain all reads in the selective sequencing scenario is therefore

$$T_{sel} = n_{t\_sel}t_{t\_seq} + n_{bg\_sel}t_{skip} + n_{ns\_sel}t_{ns} = N_{sel}\left(f_t t_{t\_seq} + f_{bg}t_{skip} + f_{ns}t_{ns}\right) \quad (3)$$

Note that this quantity represents the aggregate sum of all active pore time (not run time), effectively representing the time required for a given population of active pores to process a given population of reads at the pore occupancy/vacancy rate implied by the proportions of target, background, and non-sequence reads. For the case of no selection (subscript 0), e.g., odd control channels operating in parallel with even RUBRIC channels but sequencing everything, the total read count of the non-selecting experiment is given as in equation (1) by

$$N_0 = n_{t\_0} + n_{bg\_0} + n_{ns\_0} = f_t N_0 + f_{bg} N_0 + f_{ns} N_0 \quad (4)$$

We can likewise express the cumulative pore time for the non-selection case as:

$$T_0 = n_{t\_0} t_{t\_seq} + n_{bg\_0} t_{bg\_seq} + n_{ns\_0} t_{ns} = N_0 \left( f_t t_{t\_seq} + f_{bg} t_{bg\_seq} + f_{ns} t_{ns} \right) \quad (5)$$

For randomly fragmented target and non-target DNA from the same source, $t_{bg\_seq}$ = $t_{t\_seq}$, whereas the average sequence time for target and background fractions could be different if their size distributions were substantially different (e.g., amplicons of different lengths, fragments produced by targeted cuts, etc.). Note that the typical/average duration of non-sequence reads $t_{ns}$ within a run is presumed to be independent of selection, an approximation that is substantially supported by the empirical data and that may reflect the regular sampling interval of the Event Sampler. In practice, $t_{ns}$ is inferred from the total (odd) active pore time not occupied by fast5-producing reads divided by the count of reads without fast5s.

*Table S1: Model parameters derived for the initial set of RUBRIC real-time selective sequencing experiments.*

| Run | $T_0$ (pore-min)[1] | $T_{even}$ (pore-min)[2] | $T_{ns\_0}$ (pore-min)[3] | $n_{t\_0}$ | $n_{bg\_0}$ | $n_{ns\_0}$ | $f_t$ | $f_{bg}$ | $f_{ns}$ | $g_t$ | $g_{bg}$ | $t_{t\_seq}$ (s) | $t_{bg\_seq}$ (s) | $t_{skip}$ (s) | $t_{ns}$ (s) | Average Decision Time (s) | Enrich. Ratio (Model)[4] | Enrich. Ratio (Actual)[5] |
|---|---|---|---|---|---|---|---|---|---|---|---|---|---|---|---|---|---|---|
| A1 | 30,651 | 30,764 | 21,513 | 8,246 | 18,031 | 248,325 | 0.0300 | 0.0657 | 0.9043 | 0.3138 | 0.6862 | 21.75 | 20.46 | 3.51 | 5.20 | 2.53 | 1.199 | 1.102 |
| A2 | 29,643 | 29,163 | 19,147 | 8,140 | 19,706 | 226,973 | 0.0319 | 0.0773 | 0.8907 | 0.2923 | 0.7077 | 23.96 | 22.06 | 2.76 | 5.06 | 0.80 | 1.272 | 1.293 |
| B1 | 34,961 | 34,499 | 22,788 | 10,480 | 27,430 | 325,453 | 0.0288 | 0.0755 | 0.8957 | 0.2764 | 0.7236 | 20.54 | 18.78 | 8.68 | 4.20 | 6.71 | 1.152 | 1.028 |
| B1* | 2,341 | 2,309 | 1,662 | 663 | 1,564 | 24,463 | 0.0248 | 0.0586 | 0.9166 | 0.2977 | 0.7023 | 19.22 | 17.87 | 4.61 | 4.08 | 1.71 | 1.173 | 1.181 |
| B2 | 31,814 | 32,953 | 22,904 | 9,218 | 22,703 | 348,340 | 0.0242 | 0.0597 | 0.9161 | 0.2888 | 0.7112 | 17.90 | 16.28 | 3.70 | 3.95 | 0.79 | 1.176 | 1.212 |
| C | 40,882 | 37,263 | 26,648 | 12,281 | 27,712 | 240,803 | 0.0437 | 0.0987 | 0.8576 | 0.3071 | 0.6929 | 22.24 | 20.96 | 3.25 | 6.64 | 1.45 | 1.250 | 1.271 |
| D | 44,564 | 43,742 | 34,399 | 11,338 | 33,074 | 535,984 | 0.0195 | 0.0570 | 0.9235 | 0.2553 | 0.7447 | 15.00 | 13.30 | 3.47 | 3.85 | 0.47 | 1.138 | 1.185 |
| E1 | 25,439 | 26,327 | 20,831 | 4,062 | 12,671 | 136,724 | 0.0265 | 0.0826 | 0.8910 | 0.2428 | 0.7572 | 18.86 | 15.77 | 2.93 | 9.14 | 0.25 | 1.119 | 1.067 |
| E2 | 39,025 | 40,131 | 32,770 | 5,893 | 19,533 | 280,768 | 0.0192 | 0.0638 | 0.9170 | 0.2318 | 0.7682 | 16.64 | 14.19 | 3.41 | 7.00 | 0.29 | 1.099 | 0.920 |
| E2* | 37,783 | 38,877 | 31,701 | 5,743 | 18,988 | 271,250 | 0.0194 | 0.0642 | 0.9164 | 0.2322 | 0.7678 | 16.65 | 14.18 | 3.10 | 7.01 | 0.29 | 1.102 | 0.924 |
| F | 174,491 | 175,474 | 171,532 | 755 | 11,855 | 1,732,119 | 4.327E-04 | 6.795E-03 | 0.9928 | 0.0599 | 0.9401 | 9.84 | 14.35 | 4.00 | 5.94 | 0.23 | 1.012 | 1.056 |
| F* | 165,727 | 165,882 | 162,870 | 721 | 11,406 | 1,641,069 | 4.361E-04 | 6.899E-03 | 0.9927 | 0.0595 | 0.9405 | 9.82 | 14.41 | 2.89 | 5.95 | 0.23 | 1.013 | 1.092 |
| G | 8,999 | 8,912 | 7,041 | 74 | 16,107 | 99,229 | 6.412E-04 | 0.1396 | 0.8598 | 0.0046 | 0.9954 | 6.19 | 7.27 | 4.03 | 4.26 | 0.89 | 1.107 | 1.082 |
| G* | 6,969 | 6,924 | 5,380 | 63 | 13,195 | 78,526 | 6.864E-04 | 0.1438 | 0.8556 | 0.0048 | 0.9952 | 6.21 | 7.20 | 3.55 | 4.11 | 0.91 | 1.130 | 1.092 |

1 Integrated active pore-time for odd numbered pores
2 Integrated active pore-time for even numbered pores
3 Total (odd channel) active pore time not occupied by fast5-producing reads (i.e., empty/vacant pore time)
4 Absolute enrichment ratio predicted by equation (6) or equivalently equation (13)
5 Empirically observed enrichment (throughput) ratio = even sampled read count / odd sampled read count normalized by total even and odd active pore times, respectively.

* Dataset time-filtered to eliminate reads from periods of failed skipping, see Supplemental Section S-3.

Assuming equivalent pore-time utilized by both selecting (e.g., RUBRIC) and non-selecting (control) pores, as in the case of a substantially equal number of even and odd channels operating concurrently for an equal amount of sequencing run time, we equate $T_{sel}$ in equation (3) and $T_0$ in equation (5) to solve for the enrichment ratio:

$$\frac{N_{sel}}{N_0} = \frac{f_t t_{t\_seq} + f_{bg} t_{bg\_seq} + f_{ns} t_{ns}}{f_t t_{t\_seq} + f_{bg} t_{skip} + f_{ns} t_{ns}} \quad (6)$$

This ratio represents the maximum increase in read throughput theoretically obtainable by real-time selection for a particular library (determining $f_t$, $f_{bg}$, $f_{ns}$, $t_{t\_seq}$, $t_{bg\_seq}$, and $t_{ns}$) and computing setup (in part determining $t_{skip}$).

Because read fractions/proportions are fixed as noted above, this ratio therefore also represents the upper bound on the absolute (numerical) enrichment of target reads achievable via selection. Note that equation (6) only applies where $t_{bg\_seq} \geq t_{skip}$. If fragment-length dependent $t_{bg\_seq}$ is less than $t_{skip}$, then skipping will have no effect and equation (6) behaves as though $t_{skip} = t_{bg\_seq}$, producing a ratio of unity. For the idealized case of 100% pore occupancy, the non-sequence terms drop out, and for target and background reads with equivalent size distributions ($t_{t\_seq} = t_{bg\_seq}$), equation (6) simplifies to

$$\frac{N_{sel}}{N_0} = \frac{(f_t + f_{bg})t_{t\_seq}}{f_t t_{t\_seq} + f_{bg} t_{skip}} \qquad (7)$$

Which, in the limiting case of instantaneous skipping ($t_{skip} = 0$), further simplifies to $1 + f_{bg}/f_t$.

As equation (6) illustrates, deviation of the enrichment ratio away from unity is solely a consequence of the difference between $t_{bg\_seq}$ and $t_{skip}$ scaled by the non-target read fraction $f_{bg}$. Accordingly, the greatest benefit from selection is obtained in cases where the fraction of background reads is comparatively large and/or $t_{bg\_seq} >> t_{skip}$, corresponding to long background reads with pore transit times substantially longer than the time required for the selection process to produce and execute a skip decision. For cases involving high non-sequence read fractions (~90%) as seen in the experiments detailed here, the benefit of selection is significantly muted because the time saved by skipping only affects roughly 2/3 of 10% of all reads, yielding a predicted maximum enrichment ratio of about 1.18 for experiment B2, assuming perfect selection. Figure S8 plots the value of the enrichment ratio derived from equation (6) across a range of parameters assuming equivalent target and background size distributions, while Table S1 provides model parameters determined for the runs in this article.

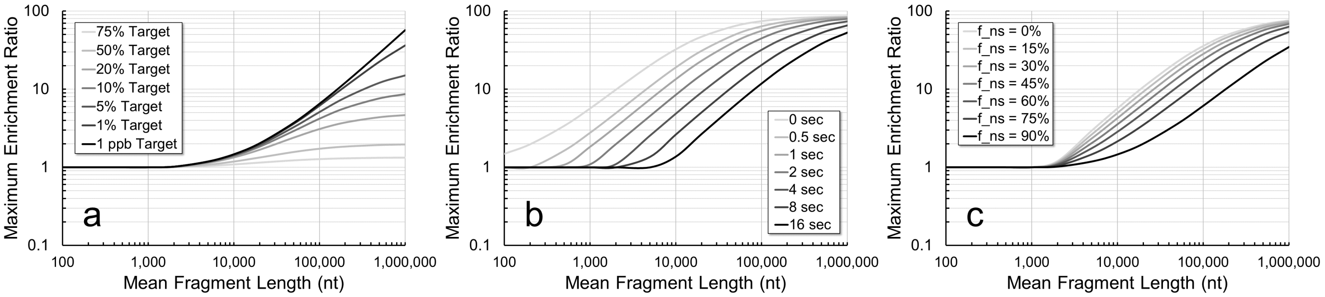


*Figure S8: Bounding limits on enrichment / throughput enhancement performance for selected library and computing parameters evaluated as in equation (6). (a) Selection for varying target:background ratios ($t_{skip}$=3.7 s, $f_{ns}$=0.90). (b) Selection for varying skip times ($f_t$=0.01, $f_{ns}$=0.1). (c) Selection for varying non-sequence read fractions ($f_t$=0.01, $t_{skip}$=3.7 s). For all plots, $t_{ns}$=4 s and fragment length-derived average sequencing times assume a 450 bases/s DNA translocation rate through the pore.*

While equation (6) is useful in analyzing existing Read Until run data, it is not ideal for predicting selection performance *a priori* or from standard MinKNOW sequence data, e.g., without the benefit having already obtained Event Sampler-reported non-sequence read counts via RUBRIC. Accordingly, we develop a modified formulation of the enrichment ratio that can be derived entirely from the output of a typical MinION sequencing run (non-selecting) performed with a representative library of interest. In the preceding analysis, we addressed Event Sampler-derived read counts (*N*, *n*) including those that did not yield fast5s. Here, we focus on MinKNOW-output fast5 counts exclusively (*M*, *n*). Analogous to equations (1) and (2) above, the total target and background counts for non-selecting and selecting runs are given by

$$M_0 = n_{t\_0} + n_{bg\_0} = M_0(g_t + g_{bg}) \ \ (8) \quad , \quad M_{sel} = n_{t\_sel} + n_{bg\_sel} = M_{sel}(g_t + g_{bg}) \ \ (9)$$

Like the read fractions (*f*) described above, sequence file fractions (*g*) are likewise presumed to be constant and independent of selection, conforming to $g_t/g_{bg} = f_t/f_{bg}$. Note that while the relevant counts of sequence-producing reads in the non-selection (odd pore) case can be derived directly from MinKNOW data, the true number of skip decision fast5s resulting from selection will not necessarily be knowable in advance due to the uncertainty about when/whether MinKNOW creates fast5s for skipped reads. As a consequence, here $n_{bg\_sel}$ is understood as the count of skip decision reads upstream of the fast5/no fast5 determination rather than the final observed count of skip decision fast5s.

Next, we express the total pore time of a non-selecting run as the sum of all target read, background read, and non-sequence or open pore time, and ultimately the average durations of target and background reads as in equation (5) above:

$$T_0 = n_{t\_0} t_{t\_seq} + n_{bg\_0} t_{bg\_seq} + T_{ns\_0} = M_0 \left( g_t t_{t\_seq} + g_{bg} t_{bg\_seq} \right) + T_{ns\_0} \quad (10)$$

Where $T_{ns\_0}$ is the total non-sequence or open pore time. Note that all these parameters can be determined empirically from a given non-selecting model sequencing experiment: target and background fast5 counts, average fragment lengths/durations, total active pore time, and aggregate pore vacancy time. For the case of selective sequencing, equation (10) becomes

$$T_{sel} = n_{t\_sel} t_{t\_seq} + n_{bg\_sel} t_{skip} + T_{ns\_sel} = M_{sel} \left( g_t t_{t\_seq} + g_{bg} t_{skip} \right) + T_{ns\_sel} \quad (11)$$

Unlike equation (10), here the non-sequence term is not obviously determined *a priori* from quantities in a non-selecting model sequencing experiment. Equating $T_0$ and $T_{sel}$ from equations (10) and (11) as in the preceding analysis and rearranging, we obtain the enrichment ratio

$$\frac{M_{sel}}{M_0} = \frac{g_t t_{t\_seq} + g_{bg} t_{bg\_seq} + T_{ns\_0}/M_0}{g_t t_{t\_seq} + g_{bg} t_{skip} + T_{ns\_sel}/M_{sel}} \quad (12)$$

Which is very similar in form to equation (6). The term, $T_{ns\_0}/M_0$, in the numerator is a constant readily obtained empirically from a model MinION run, reflecting the average amount of open pore time per fast5-producing read. While read counts (*n*) and associated time fractions will change as a consequence of selection, read and fast5 proportions (*f* and *g*) are not affected by selection. Analogous to our treatment of the non-sequence read fraction product ($f_{ns} t_{ns}$) in the derivation of equation (6), which was found to be largely consistent regardless of selection, we argue that the same should be true of the average open pore time per fast5, allowing us to equate $T_{ns\_0}/M_o$ = $T_{ns\_sel}/M_{sel}$ and revise equation (12) to our final result

$$\frac{M_{sel}}{M_0} = \frac{g_t t_{t\_seq} + g_{bg} t_{bg\_seq} + T_{ns\_0}/M_0}{g_t t_{t\_seq} + g_{bg} t_{skip} + T_{ns\_0}/M_0} \quad (13)$$

Here, the enrichment ratio is expressed solely in terms of parameters that can be derived from a sample non-selecting sequencing experiment, with the exception of $t_{skip}$, which must be assumed. In practice, empirically determined $T_{ns\_sel}/M_{sel}$ can differ significantly from $T_{ns\_0}/M_o$ due to the unpredictable effect of skipping on fast5 creation, so the assumed equality of these terms again reflects an idealized condition immediately upstream of the decision process. As in equation (6), equation (13) only applies when $t_{bg\_seq} \geq t_{skip}$. In the limiting case of 100% pore occupancy, the $T_{ns\_0}/M_o$ terms drop out, and for target and background reads with equivalent size distributions equation (13) simplifies to

$$\frac{M_{sel}}{M_0} = \frac{(g_t + g_{bg}) t_{t\_seq}}{g_t t_{t\_seq} + g_{bg} t_{skip}} \quad (14)$$

For the further idealized case of instantaneous skipping with no latency ($t_{skip} = 0$), the expression collapses to 1+ $g_{bg}/g_t$.

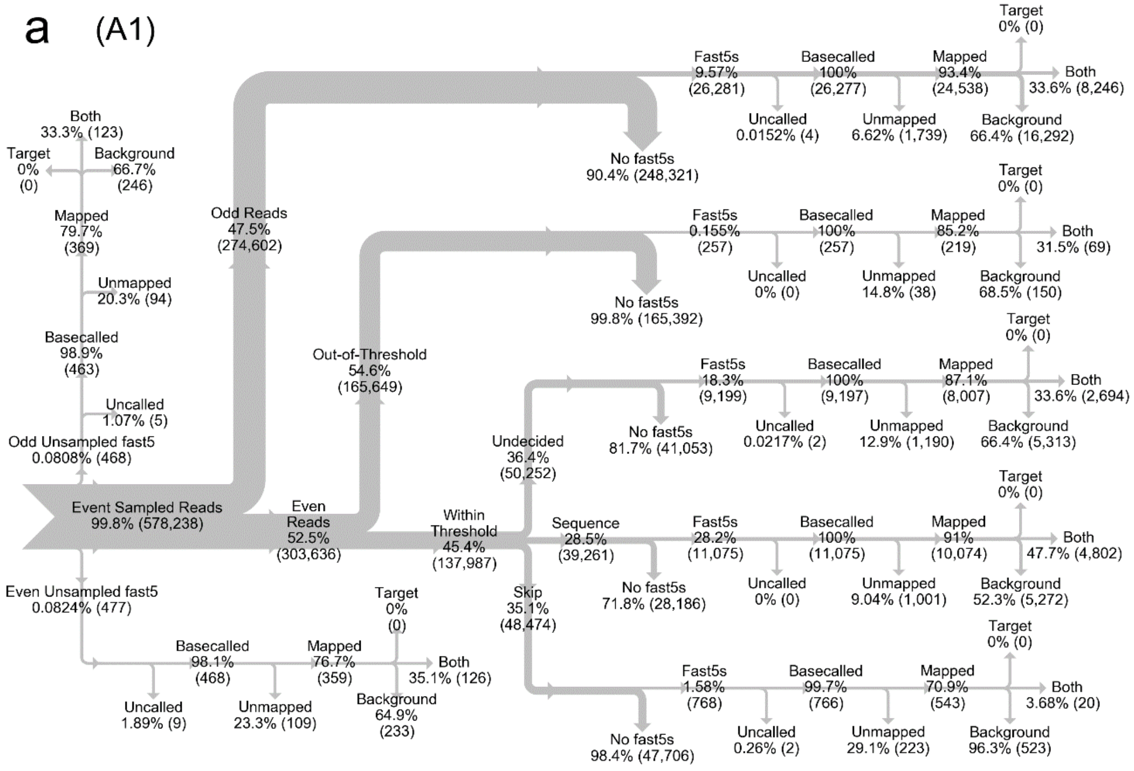


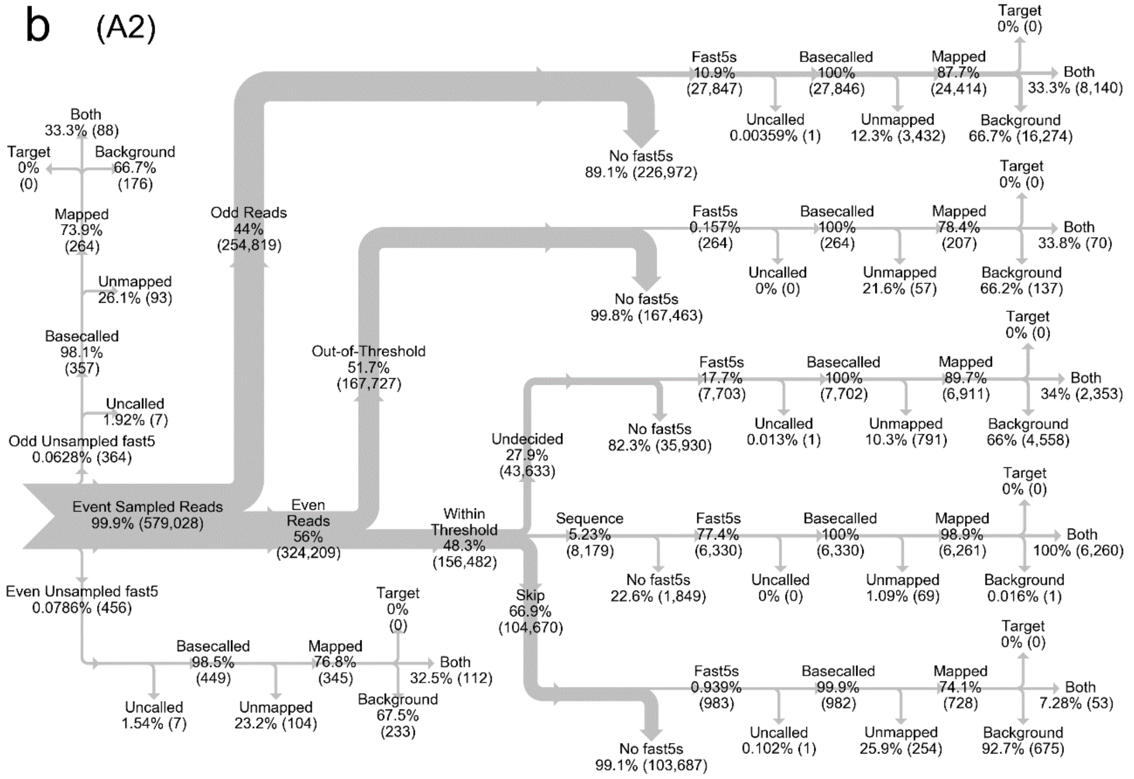


*Figure S9 (a-b): Sankey data flow diagrams for EagI-digested lambda DNA preliminary RUBRIC experiments A1-A2.*

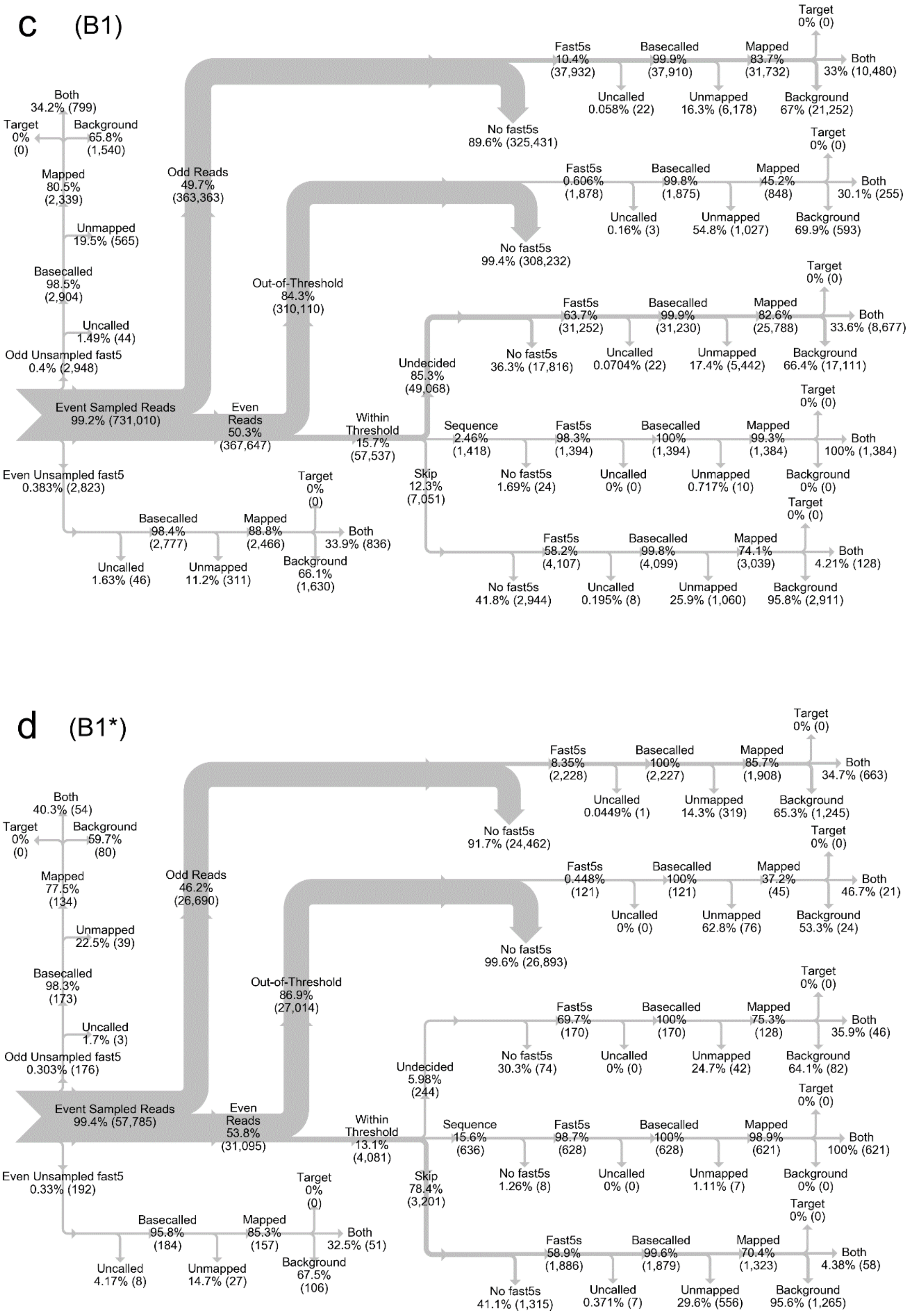


*Figure S9 (c-d): Sankey data flow diagrams for EagI-digested lambda DNA preliminary RUBRIC experiment B1 and skip-fail filtered dataset B1*.*

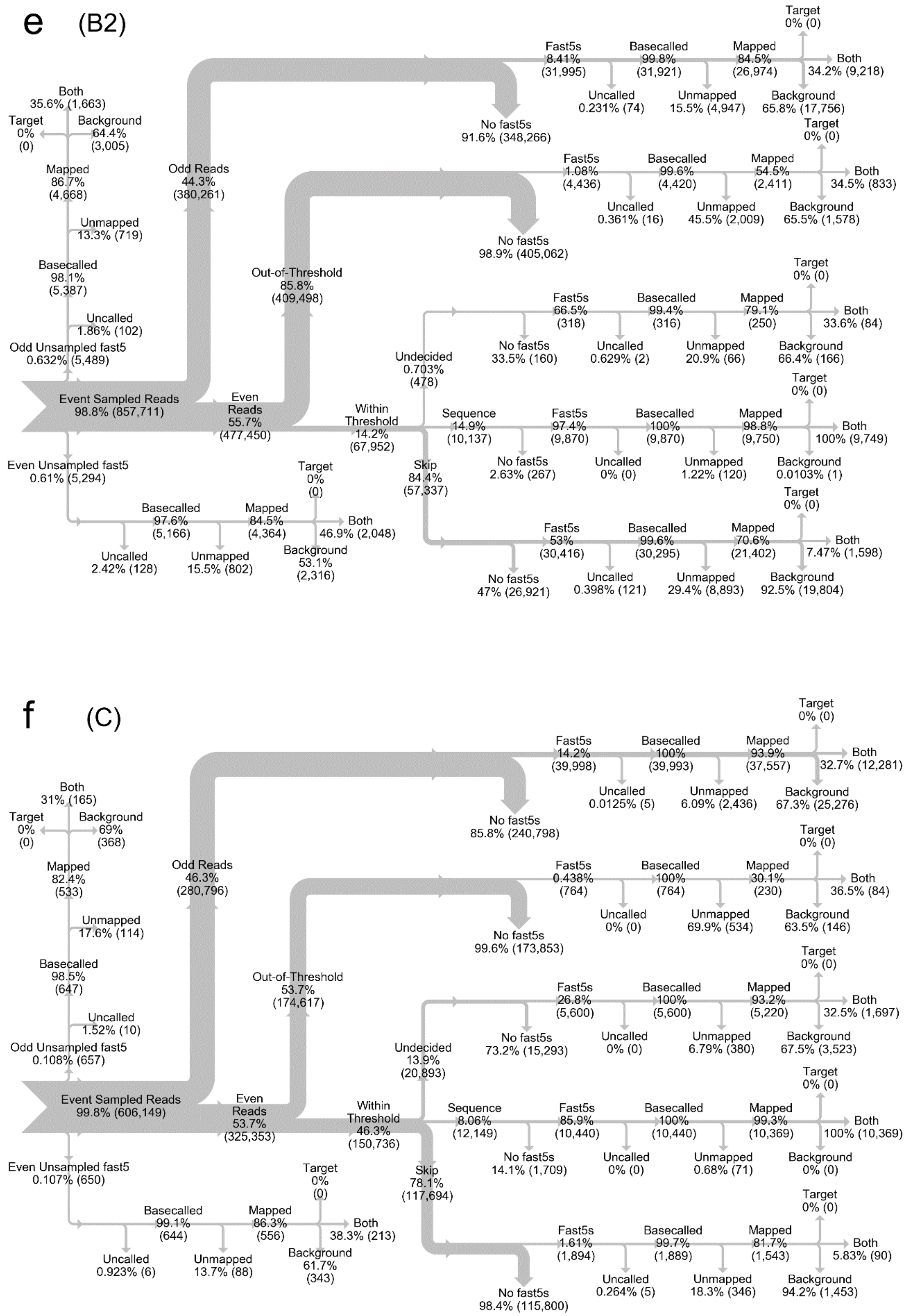


*Figure S9 (e-f): Sankey data flow diagrams for EagI-digested lambda DNA mainline RUBRIC experiments B2 and C.*

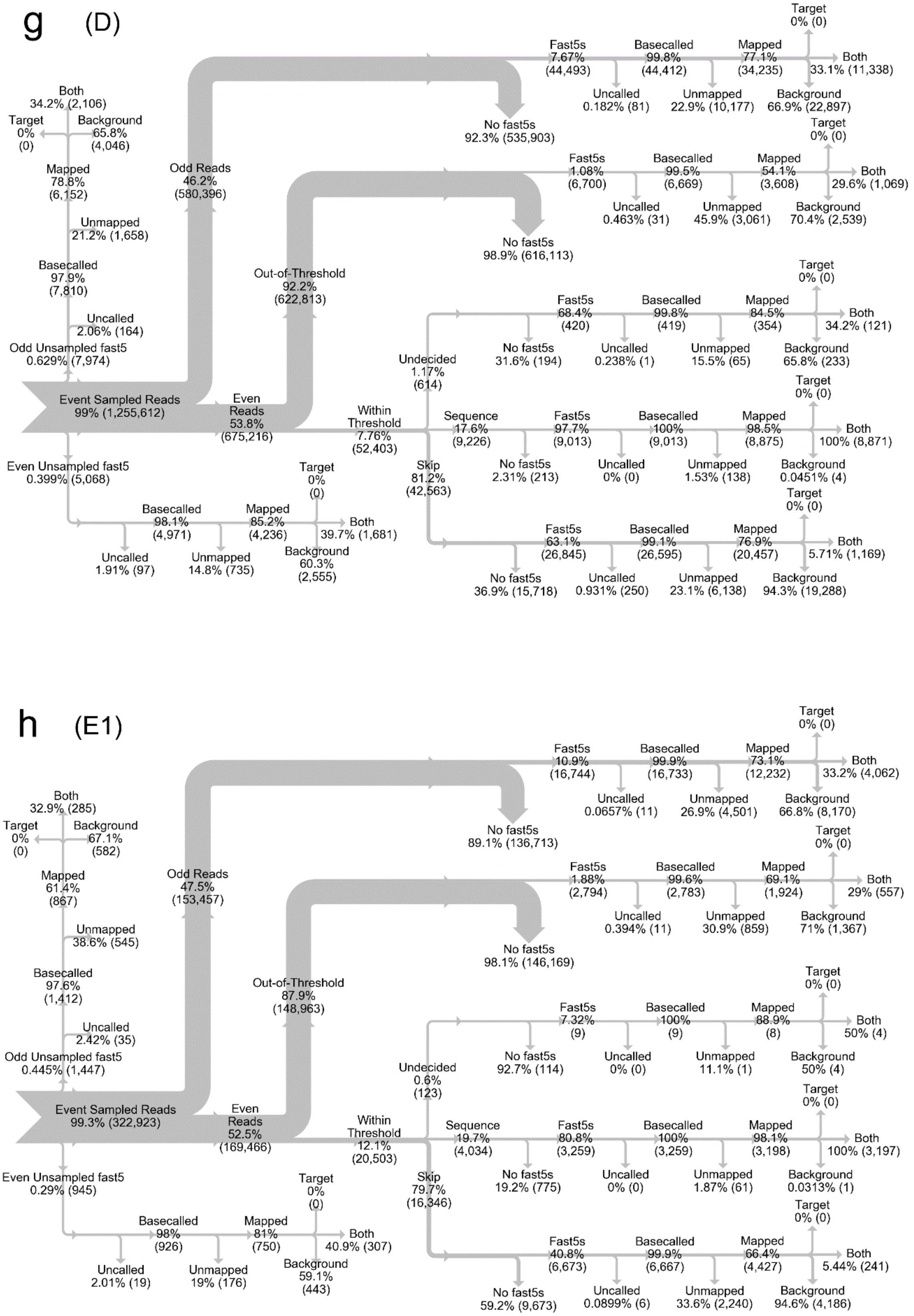


*Figure S9 (g-h): Sankey data flow diagrams for EagI-digested lambda DNA mainline RUBRIC experiments D and E1.*

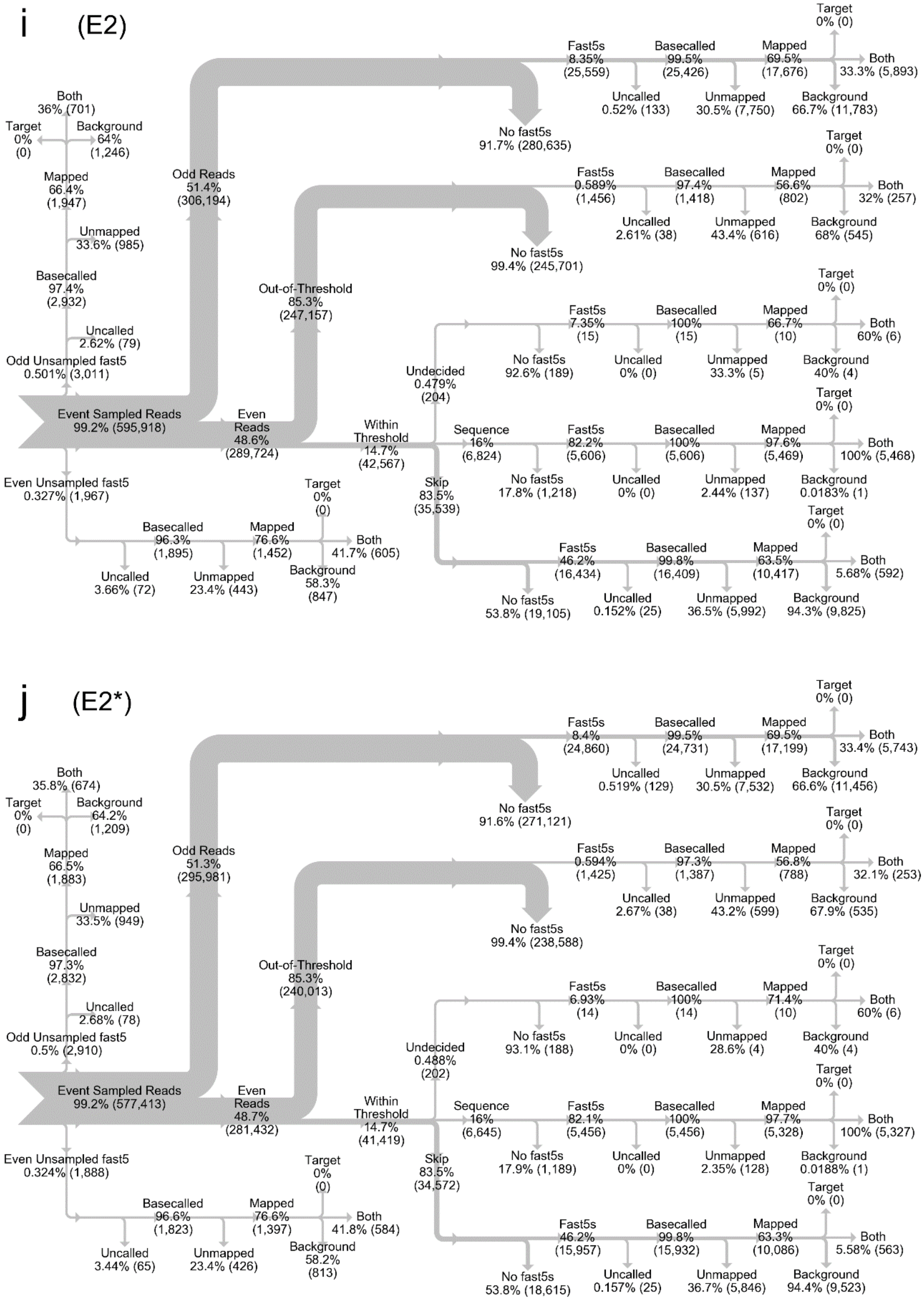


*Figure S9 (i-j): Sankey data flow diagrams for EagI-digested lambda DNA mainline RUBRIC experiment E2 and skip-fail filtered dataset E2*.*

k (F)

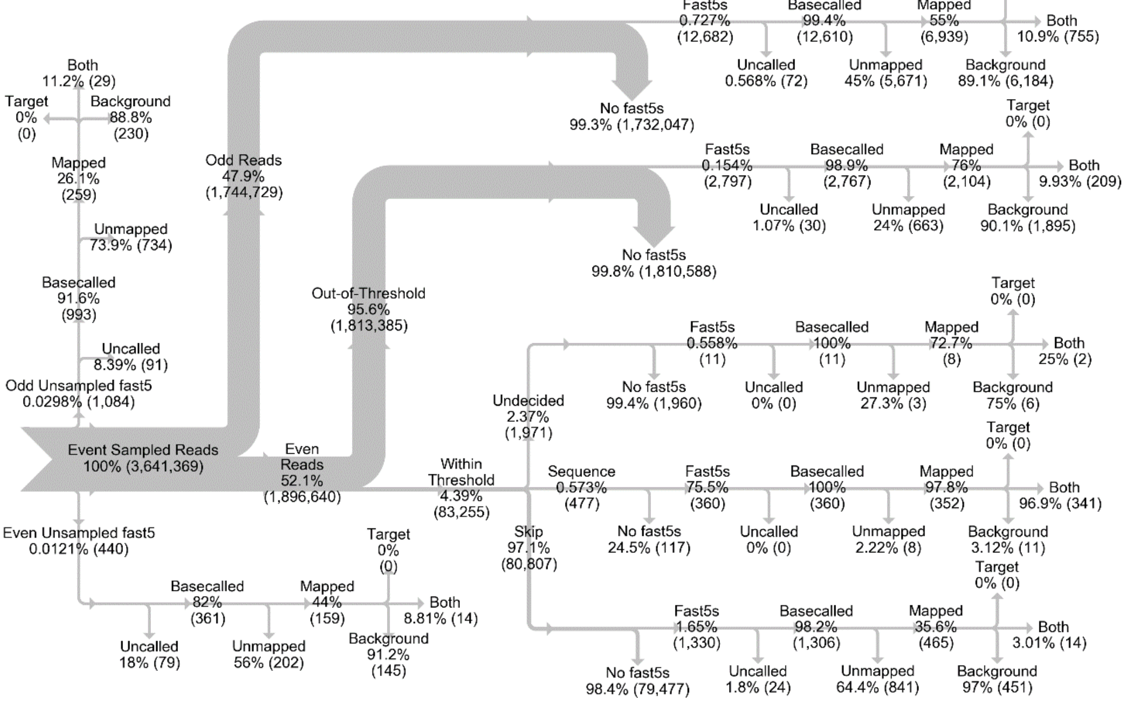


l (F*)

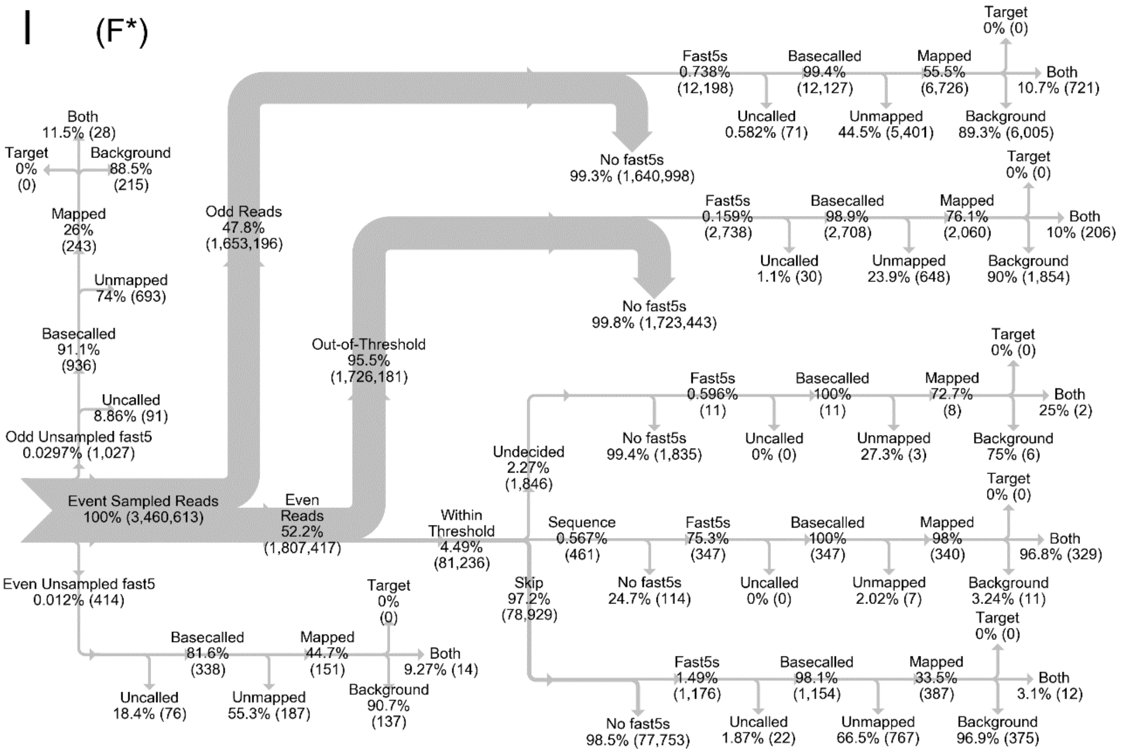


*Figure S9 (k-l): Sankey data flow diagrams for CRISPR/Cas9-excised E. coli rDNA sample use case RUBRIC experiment F and skip-fail filtered dataset F*.*

m (G)

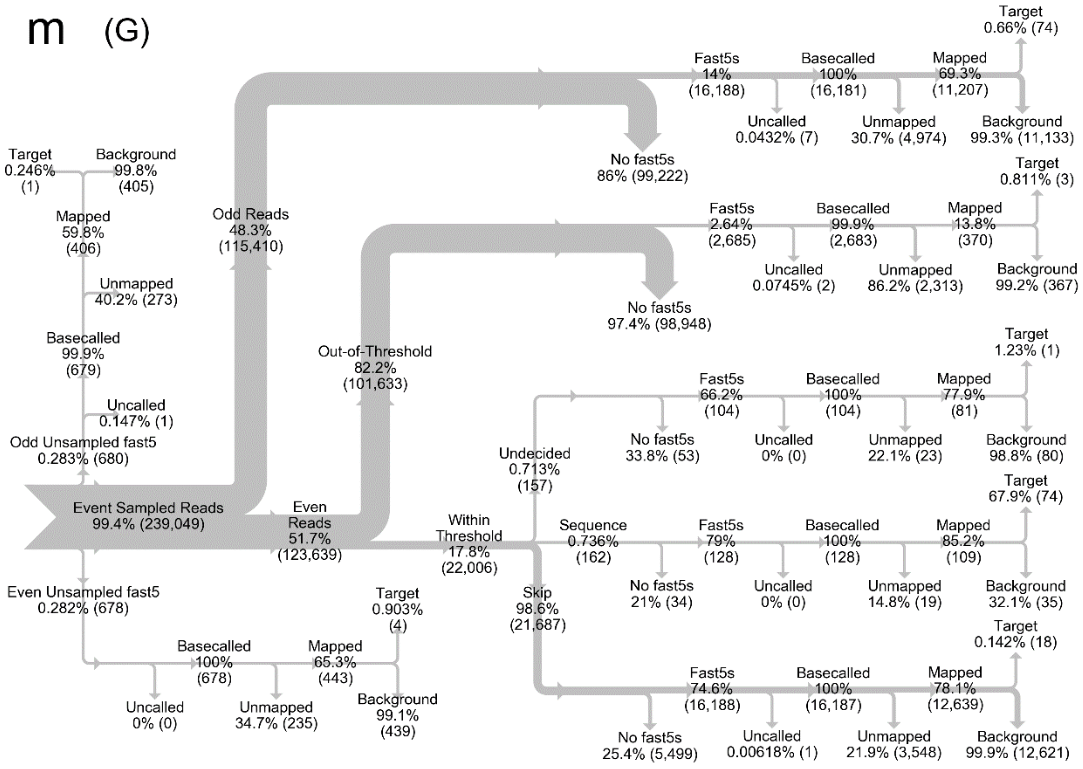


n (G*)

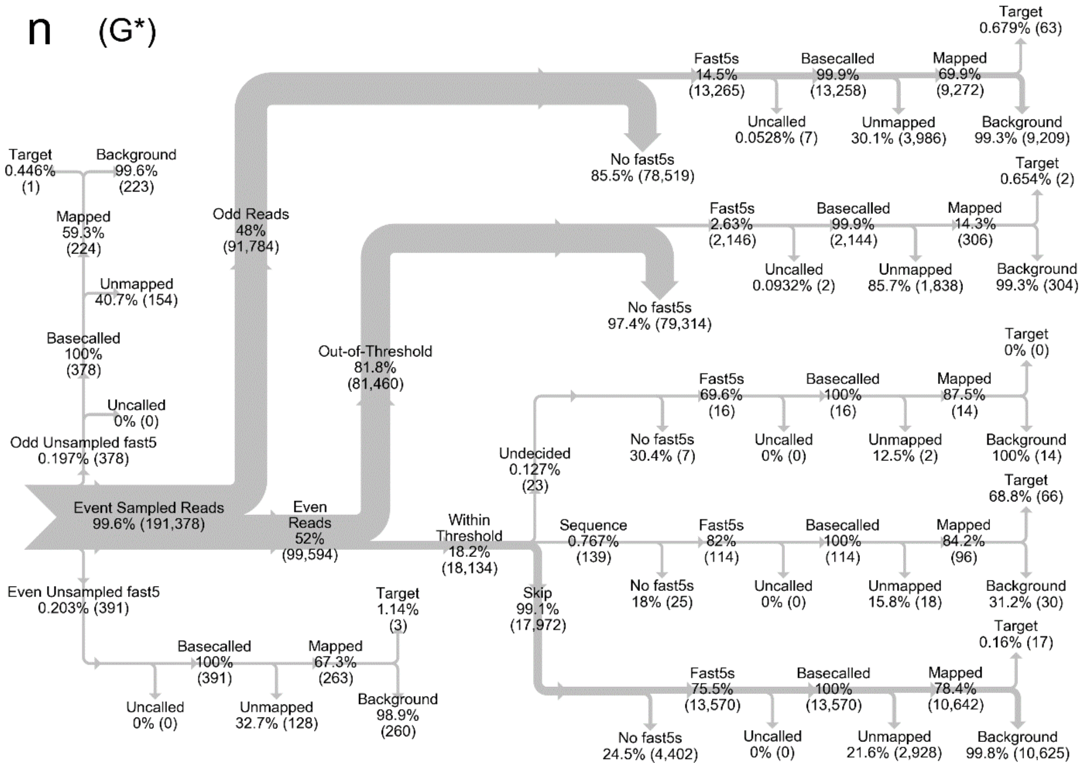


*Figure S9 (m-n): Sankey data flow diagrams for 1% E. coli / 99% human gDNA sample use case RUBRIC experiment G and skip-fail filtered dataset G*.*

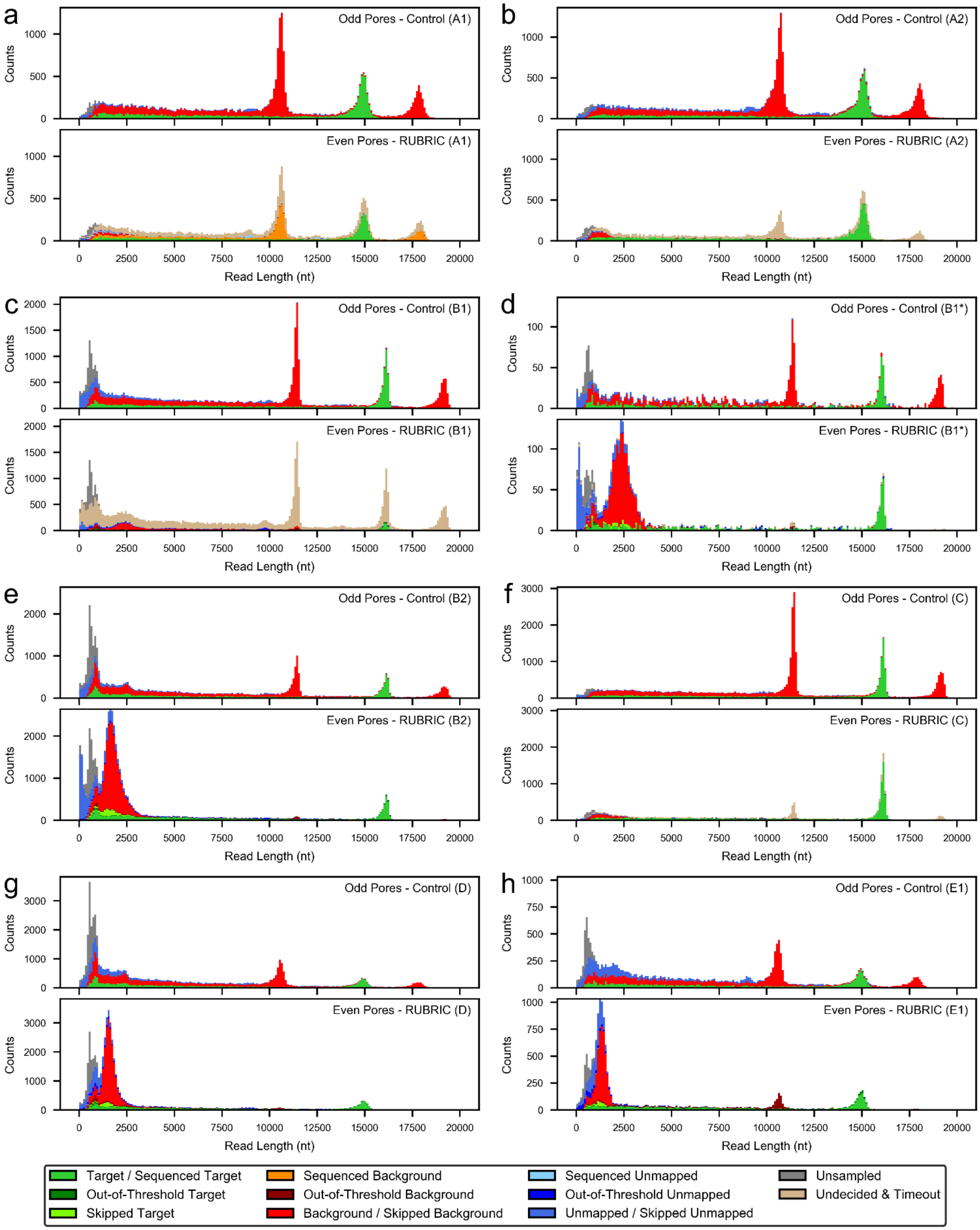


*Figure S10 (a-h): Read length histograms for lambda DNA experiments A1-E1 and filtered dataset B1* illustrating the distribution of different read types (target, non-target, unmapped) and their fate as a function of RUBRIC selection applied to even numbered pores.*

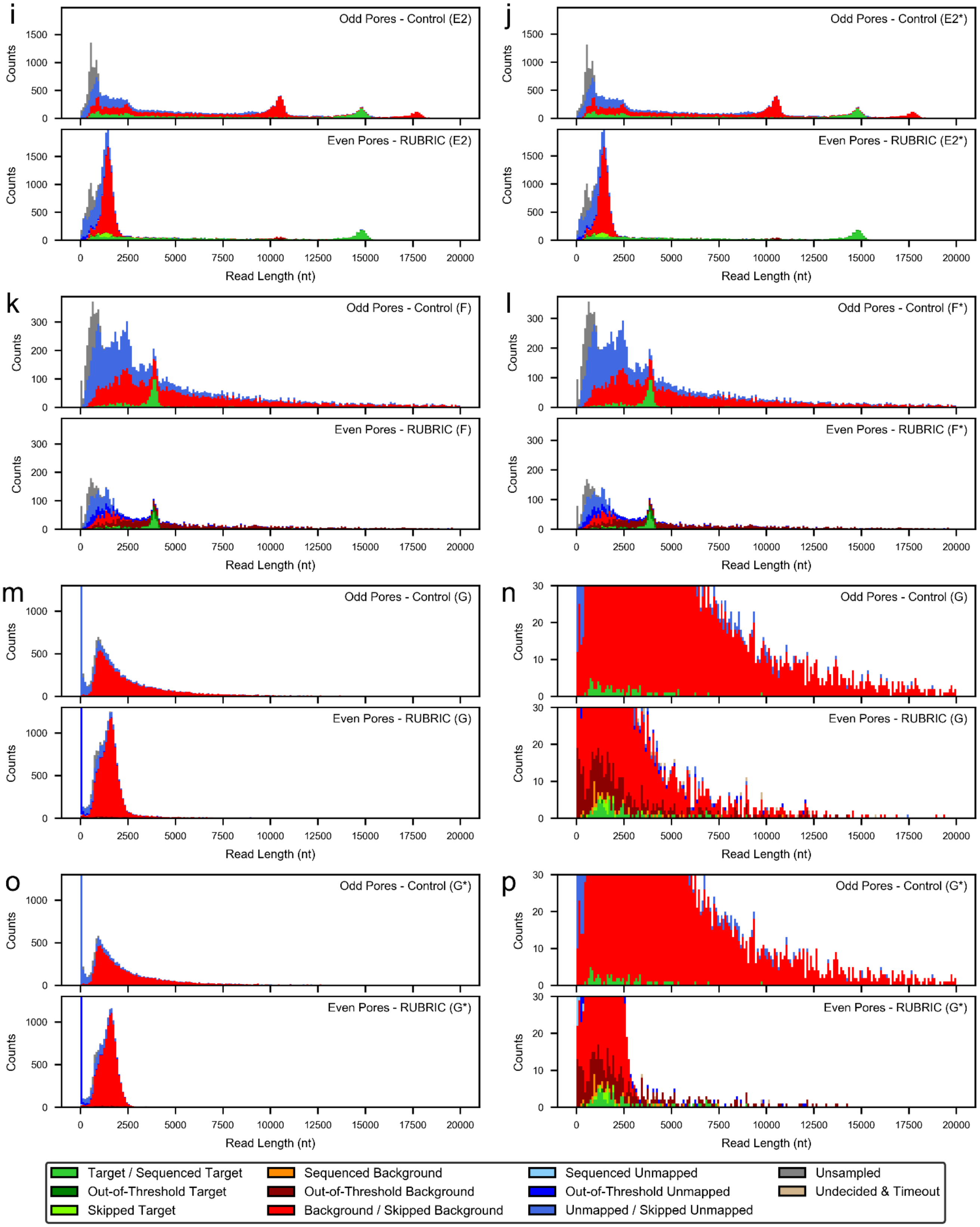


*Figure S10 (i-p): Read length histograms for lambda DNA experiment E2 and filtered dataset E2*, Cas9-exised rDNA experiment F and filtered dataset F*, and human/E. coli experiment G and filtered dataset G*, illustrating the distribution of different read types (target, non-target, unmapped) and their fate as a function of RUBRIC selection applied to even numbered pores.*
